# Canonical Wnt pathway controls mESCs self-renewal through inhibition of spontaneous differentiation via β-catenin/TCF/LEF functions

**DOI:** 10.1101/661777

**Authors:** Francesco Aulicino, Francesco Sottile, Elisa Pedone, Frederic Lluis, Lucia Marucci, Maria Pia Cosma

**Affiliations:** Centre for Genomic Regulation (CRG), Dr Aiguader 88, 08002, Barcelona, Spain; Department of Biochemistry, University of Bristol, Bristol BS8 1UB, UK; Department of Engineering Mathematics, University of Bristol, Bristol BS8 1UB, UK; Department of Development and Regeneration, Stem Cell Institute, KU Leuven, B-300 Leuven, Belgium; Universitat Pompeu Fabra (UPF), Barcelona, Spain; ICREA, Pg. Lluis Companys 23, Barcelona 08010, Spain; Guangzhou Regenerative Medicine and Health Guangdong Laboratory (GRMH-GDL), Guangzhou 510005, China; Key Laboratory of Regenerative Biology and Guangdong Provincial Key Laboratory of Stem Cells and Regenerative Medicine, Guangzhou Institutes of Biomedicine and Health, Chinese Academy of Science, Guangzhou 510530, China

## Abstract

The Wnt/β-catenin signalling pathway is a key regulator of embryonic stem cell self-renewal and differentiation. Constitutive activation of this pathway has been shown to significantly increase mouse embryonic stem cell (mESC) self-renewal and pluripotency marker expression. In this study, we generated a novel β-catenin knock-out model in mESCs by using CRISPR/Cas9 technology to delete putatively functional N-terminally truncated isoforms observed in previous knock-out models. While we showed that aberrant N-terminally truncated isoforms are not functional in mESCS, we observed that canonical Wnt signalling is not active in mESCs, as β-catenin ablation does not alter mESC transcriptional profile in LIF-enriched culture conditions; on the other hand, Wnt signalling activation represses mESC spontaneous differentiation. We also showed that transcriptionally silent β-catenin (ΔC) isoforms can rescue β-catenin knock-out self-renewal defects in mESCs, cooperating with TCF1 and LEF1 in the inhibition of mESC spontaneous differentiation in a Gsk3 dependent manner.

## Introduction

β-catenin regulates different cellular processes spanning from development to cancer progression. In addition to its central role in *adherens* junctions, β-catenin is the key effector of the canonical Wnt signalling pathway. Exposure to canonical Wnt ligands, such as Wnt3a or small-molecule inhibitors of Gsk3 activity, triggers β-catenin stabilization and its nuclear translocation. In the nucleus, β-catenin acts as a scaffolding protein for transcriptional co-factors such as TCF/LEF family members thereby activating the expression of Wnt target genes. Over the past years, an accumulating number of evidences highlighted a key role for Wnt/β-catenin signalling in sustaining self-renewal, pluripotency, and cell-cycle progression of mouse embryonic stem cells (mESCs) [1-3] and in regulating somatic cell reprogramming [4-7]. Despite its importance in mESC physiology, β-catenin knock-out mESC lines developed so far show no defects in self-renewal or pluripotency marker expression when cultured in a medium containing serum plus the leukemia inhibitory factor (LIF), but display a strict LIF requirement for pluripotency maintenance and fail to efficiently differentiate *in vitro* [8, 9]. Surprisingly, LIF-dependency can be rescued by transcriptionally defective β-catenin isoforms (ΔC mutants), challenging the hypothesis that β-catenin transcriptional activity could be relevant for mESC pluripotency and self-renewal [8, 9]. More recently however, it has been demonstrated that the most widely used inducible β-catenin knock-out alleles lead to the production of uncharacterized N-terminally truncated β-catenin isoforms (ΔN-β catenin) during pre-implantation embryo development [10]. Here, we confirmed the production of ΔN-βcatenin isoforms also in mESCs and generated a novel β-catenin full knock-out model in mESCs using CRISPR/Cas9. We found that complete β-catenin deletion produces similar phenotypes observed in the previously described knock-out models retaining ΔN-β catenin fragments, suggesting that N-terminally truncated isoforms are biologically inactive. We furthermore analysed the impact of β-catenin loss at transcriptomic level, in presence or absence of Gsk3 chemical inhibition. Our results show that the Wnt/β-catenin pathway is not transcriptionally active in mESCs cultured in Serum/LIF. However, upon Gsk3 inhibition, we observed β-catenin dependent inhibition of differentiation markers, while the expression of pluripotency genes remained unchanged. Finally we showed that ΔC β-catenin isoforms are not entirely transcriptionally silent and their nuclear function depends on TCF/LEF factors, indeed ΔC β-catenin can inhibit mESC differentiation in absence of LIF through TCF1/LEF1 activity.

## Results

### Inducible ß-catenin knock-out alleles generate N-terminally truncated isoforms in mESCs

ß-catenin (Ctnnb1) mRNA includes 15 exons, with an Open Reading Frame (ORF) spanning from exon 2 to exon 15. Previously reported ß-catenin knock-out models in mouse embryonic stem cells (mESCs) were generated after CRE-mediated excision of a DNA fragment flanked by two LoxP sites encompassing exons 3-6 (*Ctnnb1*^*tm4Wbm*^) or 2-6 (*Ctnnb1*^*tm2Kem*^) or [11, 12]. Three groups have independently studied both alleles in mESCs and concluded that β-catenin is dispensable for mESC self-renewal [8, 9, 13].

A recent study reported that, during pre-implantation embryo development, both these ß-catenin knock-out alleles are flawed by the production of N-terminally truncated (ΔN) proteins, possibly generated by alternative splicing (*Ctnnb1*^*tm4Wbm*^) or by a secondary ATG within a Kozak consensus sequence downstream of the excised region (*Ctnnb1*^*tm2Kem*^) (**Figure 1A**) [10, 14]. Remarkably the production of these ΔN ß-catenin isoforms in mESCs has not been previously reported [8, 9, 15, 16]. We therefore tested the expression of ΔN ß-catenin isoforms by using an antibody raised against the C-terminal portion of ß-catenin in protein extracts of NLC1 and SR18 mESCs, which harbour the *Ctnnb1*^*tm4Wbm fl/fl*^ and *Ctnnb1*^*tm2Kem fl/del*^ alleles, respectively, and stably express the CRE-ERT2 construct as in [13]. Full-length ß-catenin was successfully excised upon 4-Hydroxytamoxyfene (4OHT) treatment in both cell lines, and ΔN ß-catenin isoforms with a molecular weight of approximately 48 and 52 KDa respectively were detected upon CRE-recombination as previously reported [10] (**Figure 1B**). Immunofluorescence staining did not show any clear sub-cellular localization of ΔN isoforms, which instead appeared distributed among cytoplasm and nuclei showing little or no membrane localization (**Figure 1C**).

**Figure 1.**
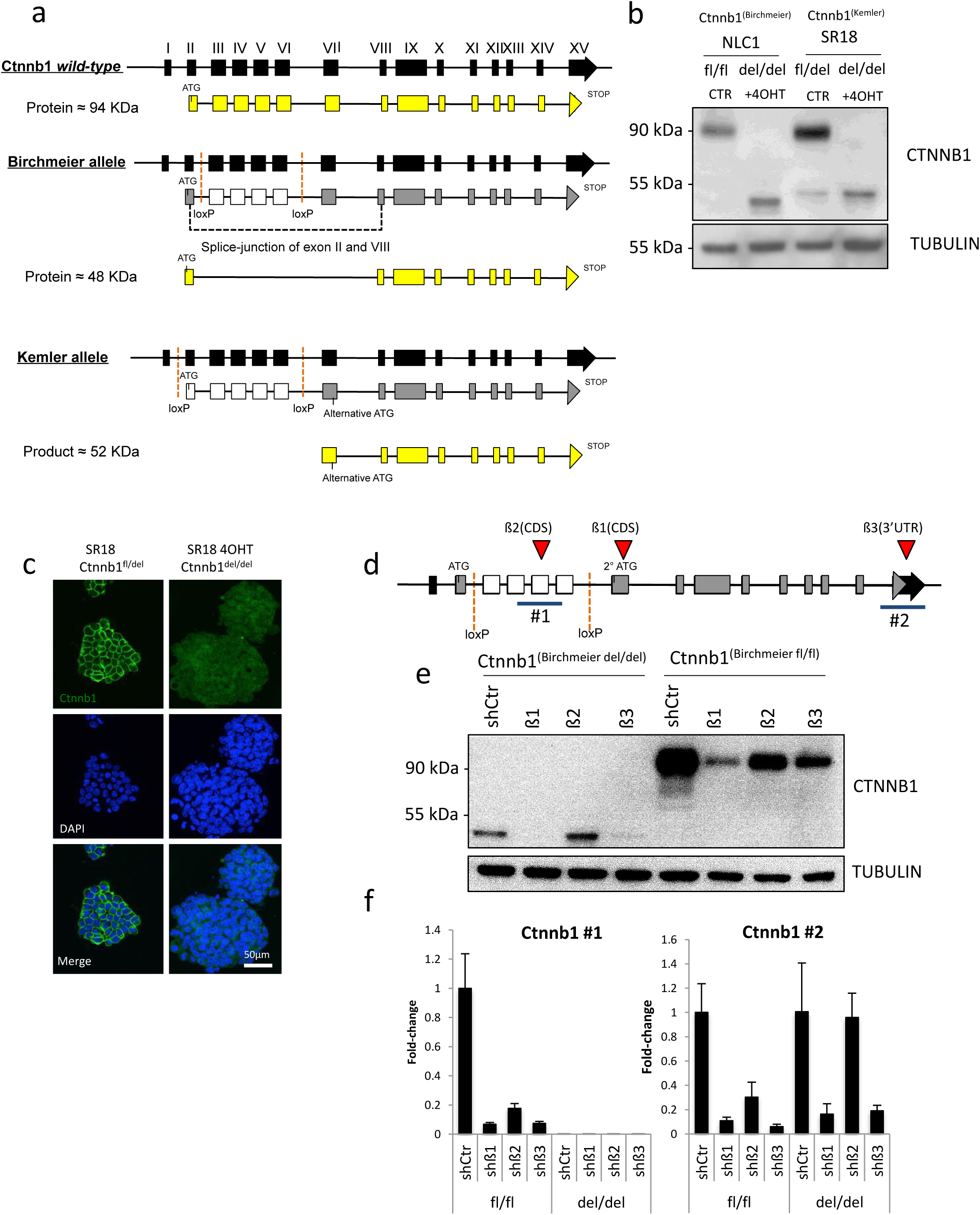
Inducible β-catenin knock-out alleles produces N-terminally truncated isoforms in mESCs. **a)** Schematic representation of murine β-catenin (Ctnnb1) locus and the two loxP alleles used for β-catenin studies in mESCs. Black boxes represent exons, yellow boxes coding exons, dashed red lines indicates loxP sites, white boxes represent exons excised upon CRE-mediated recombination of loxP sites. **b)** Western blot of NLC1 and SR18 cell lines upon 72hrs 4’-Hydroxytamoxifen treatment (+4OHT) and respective untreated controls (CTR). SR18 untreated cell line is heterozygous for full-length β-catenin deletion. **c)** β-catenin immunofluorescence staining on fixed SR18 (Ctnnb1^fl/del^) parental cell line (left) or upon 72 hours 4’-Hydroxytamoxifen treatment (right panel). DAPI was used to counterstain nuclei. A primary antibody raised against the C-terminal portion of β-catenin was used. **d)** Schematic representation of short-hairpin targeted regions (red triangles, β1, β2 and β3) and qRT-PCR amplicons (blue lines, #1 and #2) along the Ctnnb1^Birchmeier^ allele. **e)** Western blot of β-catenin of mESCs harbouring the Birchmeier β-catenin allele after (Ctnnb1^del/del^ (left)) or before (Ctnnb1^fl/fl^ (right)) CRE-mediated recombination of the lox-P sites. Cells were transduced with a control short-hairpin (shCtr) or three different short hairpins against β-catenin mRNA (β1, β2 or β3). **f)** qRT-PCR on total mRNA extracts of Ctnnb1^fl/fl^ or Ctnnb1^del/del^ cells transduced with the short-hairpin constructs used in Figure 1d. Two different amplicons were amplified to monitor deleted region (Ctnnb1 #1) or 3’UTR region (Ctnnb1 #3). GAPDH was used as housekeeping control. Error bars represents standard deviation of technical triplicates.

To assess if N-terminally truncated isoforms were a product of the Ctnnb1 genomic locus and not technical artefacts, we designed three different short hairpins against three different regions of ß-catenin mRNA (ß1-3) (**Figure 1D**) to silence ß-catenin in wild-type (*Ctnnb1*^*tm4Wbm fl/fl*^) and knock-out (*Ctnnb1*^*tm4Wbm del/del*^) mESCs. All the short hairpins successfully induced knock-down of ß-catenin at protein level (**Figure 1E**) in wild-type cells, while, as expected, N-terminally truncated isoforms were depleted efficiently by ß1 and ß3 but not by ß2 which targets the portion of mRNA excised upon CRE-mediated recombination. The same results were confirmed analysing ß-catenin mRNA levels by qRT-PCR using two different primer pairs targeting the CRE-excised region (**Figure 1F, left panel**), and the 3’UTR (**Figure 1F, right panel**). The oligonucleotides targeting the excised region failed to detect ß-catenin mRNA in knock-out cells, while the ones designed on the 3’UTR revealed that mRNA regions downstream of the excision had comparable expression levels in wild-type (shSCR fl/fl) and knock-out (shSCR del/del) cells and were efficiently silenced only by ß1 and ß3 but not, as expected, by ß2 (**Figure 1F, right panel**). All together these data demonstrate that N-terminally truncated isoforms are specifically expressed from the Ctnnb1 locus upon CRE-mediated recombination of the *Ctnnb1*^*tm2Kem*^ or the *Ctnnb1*^*tm4Wbm*^ inducible alleles as previously reported [10, 14]. Importantly, we never observed ΔN isoforms in wild-type cells, including cells expressing short-hairpins targeting ß-catenin, indicating that they are only produced upon recombination of Ctnnb1 locus.

Next, we characterized SR18 cells (*Ctnnb1*^*tm2Kem fl/del*^) upon ß-catenin CRE-mediated deletion. As previously reported [13], upon 4OHT treatment and consequent ß-catenin deletion, clone morphology and alkaline phosphatase expression (**Figure S1A**), as well as Nanog and Oct4 expression patterns, remained unaltered upon ß-catenin loss (**Figure S1B**). The lack of morphological changes upon ß-catenin deletion was likely due to compensatory effects by the up-regulation of Plakoglobin (γ-catenin) (**Figure S1C and S1D**), as previously reported [8, 9]. Furthermore, no changes were detected in the protein levels of pluripotency markers (Nanog, Oct4, Sox2) (**Figure S1D**). Although these results show that ß-catenin loss does not affect self-renewal, morphology and pluripotency markers, we asked whether the ΔN isoforms, produced by *Ctnnb1*^*tm2Kem*^ or the *Ctnnb1*^*tm4Wbm*^ alleles, could be responsible for the observed mESC phenotype.

### Generation of a new ß-catenin knock-out model in mESCs using CRISPR/Cas9 technology

We generated a full ß-catenin knock-out mESC line via CRISPR/Cas9 technology. As the presence of ΔN ß-catenin isoforms could be due either to an alternative splicing or to the presence of a secondary ATG on Exon VII (Figure 1A), we designed two different sgRNA pairwise combinations (sgRNA1+sgRNA3 and sgRNA2+sgRNA3, resulting in 5287 and 4970 bp deletion, respectively, **Figure S1E**) to induce deletions spanning both alternative splicing sites and the entire Exon VII. While the editing efficiency of sgRNA2+sgRNA3 was low, sgRNA1+sgRNA3 induced the deletion in a high percentage of cells (**Figure S1F**) with a significant loss of full-length ß-catenin protein in the pool of transfected cells (**Figure S1G and S1H**). The deletion generated by sgRNA1+sgRNA3 resulted in the production of a new ΔN-isoform with a lower molecular weight (40 KDa) with respect to the N-terminally truncated isoforms produced by *Ctnnb1*^*tm2Kem*^ (48 KDa) or the *Ctnnb1*^*tm4Wbm*^ (52 KDa) alleles (**Figure S1G**). The generation of this new ΔN isoform could only be explained by the presence of alternative downstream ATG sites, which however were not predictable *a priori*.

In order to eliminate any possible undesired gene product upon gene editing we sought to induce a 10 Kb deletion encompassing the whole gene body. For this aim, we used an additional couple of sgRNAs (sgRNA4+sgRNA5), targeting the upmost sequence before the canonical ATG and a portion of the last coding Exon (XV) (**Figure 2A**). Successful gene editing was confirmed at pool levels by PCR genotyping (**Figure S2A**). The pool of transfected cells did not show any expansion defect or cell-detachment. Single-cell clones were picked, expanded and finally three independent clones were isolated with homozygous deletions, Eß11, Eß15 and Eß47. All three clones carried homozygous deletion of a 9,7 Kb region as assessed by PCR genotyping and Sanger sequencing (**Figure 2B, S2B and Supplementary information SI1**) encompassing the whole β-catenin CDS, therefore preventing Ctnnb1 locus re-arrangement.

**Figure 2.**
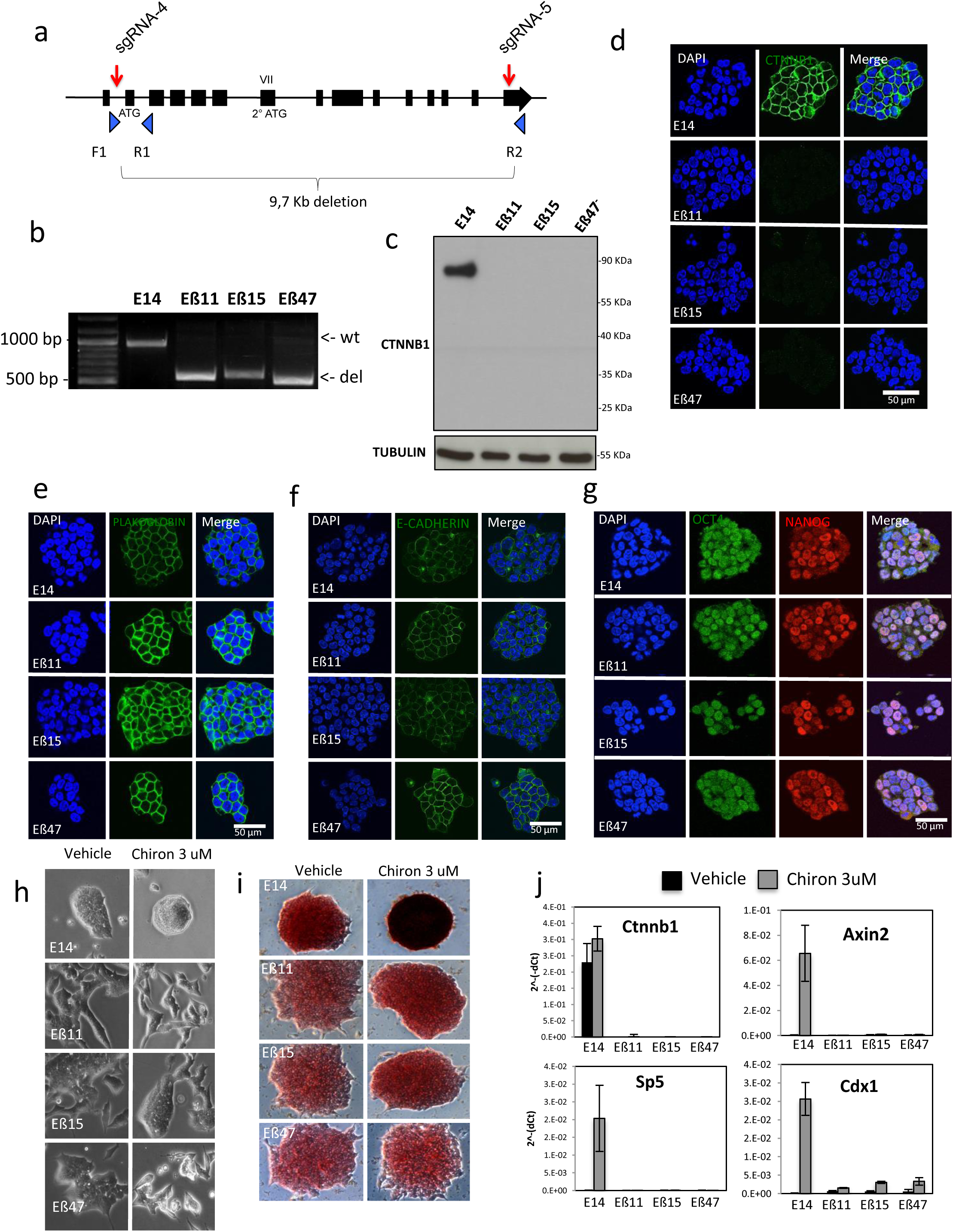
CRISPR/Cas9-mediated excision of whole Ctnnb1 locus results in a complete β-catenin knock-model in mESCs. **a)** Schematic representation of sgRNAs design for CRISPR/Cas9-mediated excision of whole β-catenin coding sequence. Red arrows indicate sgRNAs target sites. Blue triangles indicate position and orientation of oligonucleotides used for PCR genotyping. **b)** PCR genotyping of three homozigous β-catenin knock-out clones (Eβ11, Eβ15 and Eβ47) and parental E14 mESCs cells. Expected amplicon size is 951 bp for wild-type (wt) alleles and 551 bp for knock-out alleles (del). **c)** Western blot of total protein extracts from Eβ11, Eβ15, Eβ47 and wild-type E14 cells. Protein extracts were probed for β-catenin (using a C-terminally raised antibody), stripped and re-probed for Tubulin as loading control. **d-g)** Immunofluorescence in fixed parental E14, Eβ11, Eβ15 and Eβ47 cells for β-catenin (d), Plakoglobin (e), E-cadherin (f), Oct4 and Nanog (g). DAPI was used to counterstain nuclei. Scalebar= 50 μm. **h)** Phase contrast pictures of Eβ11, Eβ15, Eβ47 and parental E14 clones upon 72 hours Vehicle (0.3 % DMSO, left) or Chiron 3 μM treatment (right) in Serum/LIF. Cells were seeded at 4×10^^5^ cells/well density in multi-6 well plates. **i)** Phase contrast pictures of alkaline phosphatase staining on E14, Eβ11, Eβ15 and Eβ47 cells cultured in Serum/LIF in presence of Vehicle (0.3 % DMSO, left) or Chiron 3 μM (right) for 5 days. 300 cells were seeded in each well of a multi-6 wells plate. **j)** qRT-PCR on total RNA extracts of E14, Eβ11, Eβ15 and Eβ47 cells exposed to Vehicle (0.4% DMSO, black bars) or Chiron 3 μM (grey bars) for 72 hours. Error bars = standard deviation of technical duplicates.

Western blot analysis with an antibody raised against the C-terminal portion of ß-catenin revealed the absence of any ΔN isoform in all the analysed clones (**Figure 2C**); this, together with the absence of any detectable signal by immunofluorescence (**Figure 2D**), confirmed the complete ß-catenin depletion in the newly generated clones.

These data demonstrate that, regardless the gene-editing technology used (CRE-mediated excision or CRISPR/Cas9), gene editing by-product cannot be easily predicted, and that complete locus ablation is a viable alternative to avoid undesired protein rearrangements.

### Complete ß-catenin loss does not affect self-renewal and pluripotency markers expression in mESCs cultured in Serum/LIF

In order to characterize the impact of complete ß-catenin ablation in mESCs, we analysed the newly generated cell lines for proliferation, self-renewal and pluripotency marker expression defects under Serum/LIF culturing conditions. ß-catenin is normally found at the plasma membrane, where it physically interacts with E-cadherin and α-catenin connecting *adherens* junctions to the actin cortex. Eß11, Eß15 and Eß47 clones however did not show major morphological defects. Plakoglobin levels were upregulated in response to ß-catenin loss (**Figure 2E, S2C** and **S2D**) while E-cadherin localization and expression levels remained unchanged (**Figure 2F** and **S2E)**. We did not detect any difference in the cell-cycle (**Figure S2F**) or proliferation rate (**Figure S2G**) in all three knock-out clones with respect to the parental cell lines, which displayed overall comparable populations doubling times (**Figure S2H**).

Similarly to previously reported knock-out models [8, 9, 13], complete ß-catenin loss did not show any additional defects of Nanog and Oct4 pluripotency marker expression as their localization (**Figure 2G**) and expression levels (**Figure S2C** and **S2D**) remained similar to the parental E14 cell line in all the analysed clones.

The addition of small-molecule inhibitors of Gsk3 activity has been reported to promote mESC pluripotency and self-renewal, while inhibiting spontaneous differentiation, by increasing ß-catenin levels and activating the expression of Wnt-target genes [1].

We therefore monitored the morphological and transcriptional response of ß-catenin knock-out clones to CHIR99201 (Chiron), a selective small-molecule inhibitor of Gsk3. Wild-type and knock-out cells were cultured in Serum/LIF supplemented with Chiron or Vehicle (DMSO 0.3 ul/ml). After 72 hours of Chiron treatment, wild-type cells acquired an homogeneous round-shaped morphology with tight colony boundaries, while Eß11, Eß15 and Eß47 mESCs did not show any major morphological change with respect to Vehicle (**Figure 2H**), indicating that morphological changes induced by Gsk3-inhibition are mediated by ß-catenin.

Morphological defects were only evident when cells were reaching >50% confluency (**Figure 2H, vehicle**), while ß-catenin knock-out clones were indistinguishable from wild-type E14 if plated at clonal density (**Figure 2I, vehicle**).

These data suggest that ΔN ß-catenin isoforms are biologically inactive as complete ß-catenin depletion does not result in any additional phenotypic defect with respect to previously characterized knock-out models.

Next, we assessed the colony formation capacity and alkaline phosphatase (AP) expression of wild-type and ß-catenin knock-out cells in response to Gsk3 inhibition. Cells were plated at clonal density and while cultured in Serum/LIF in presence of Vehicle, wild-type E14 and ß-catenin knock-out clones were all positive for alkaline phosphatase (AP) expression, although ß-catenin knock-out cells displayed a slightly lower intensity (**Figure 2I, S2I** and **S2J)**. In presence of Chiron however, wild-type cells dramatically increased AP staining intensity while ß-catenin knock-out clones showed little increase in AP staining intensity (**Figure 2I, S2I** and **S2J**).

We then assessed the transcriptional response to Gsk3-inhibition in wild-type and ß-catenin knock-out clones upon 72 hrs of Chiron treatment through qRT-PCR. As expected *ß-catenin* mRNA levels were undetectable in all knock-out clones and remained unchanged upon Chiron treatment in wild-type cells. Instead, canonical Wnt targets such as *Axin2* and *Sp5* showed comparable levels between wild-type and knock-out cells while their expression was activated only in wild-type in response to Chiron treatment (**Figure 2J**). Of note, slightly higher levels of *Cdx1* mRNA were found in all knock-out clones in basal conditions with respect to wild-type cells. In addition, although the absence of β-catenin severely impaired *Cdx1* upregulation upon Chiron treatment, it did not completely abrogate it, suggesting that downstream targets of Gsk3 contribute to partially regulate its expression (**Figure 2J**).

### β-catenin ablation promotes a weak upregulation of primitive endoderm genes

According to previously published works, the canonical Wnt pathway is active in mESCs [2] and β-catenin nuclear translocation reinforces the pluripotency network by upregulating pluripotency genes such as Nanog, Esrrb or Tcfp2l1 [17-19] mainly by inhibiting Tcf3 repressive activity on their promoters [18]. Based on this model, a reasonable consequence of β-catenin ablation would be the increase in Tcf3 repressive activity on core-pluripotency network genes with an expected destabilization of self-renewal and pluripotency features. Complete β-catenin loss did not however affect Tcf3 levels or its subcellular localization (**Figure S2K**), suggesting that in absence of exogenous Wnt3a or Gsk3 inhibitors, β-catenin does not control Tcf3 activity. On the other hand, previous β-catenin knock-out model did not reveal major transcriptomic defects [8] but, due to the lack of characterization of truncated isoforms, a possible explanation for this apparently controversial phenotype could be due to ΔN β-catenin isoforms retaining some of the functions of the full-length, as recently proposed in pre-implantation embryo development [10].

We therefore sought to analyse the expression profile of E14 parental cell line and the Eβ47 knock-out clone through high-throughput RNA-seq in cells cultured in serum/LIF culturing conditions upon treatment with either vehicle (0.3% DMSO) or 3 μM Chiron for 72 hours (**Figure 3A**). Since the impact of complete β-catenin loss in mESCs has never been assessed before at transcriptomic level, we designed an assay to study, through different comparisons between samples, the β-catenin and Gsk3 dependent transcriptional changes in mESCs (**Figure 3A**). As expected, β-catenin was not detectable at transcriptional level in Eβ47 clone (**Figure S3A**), confirming the difference with previous KO models in which *β-catenin* mRNA was still detectable (**Figure 1F**).

**Figure 3.**
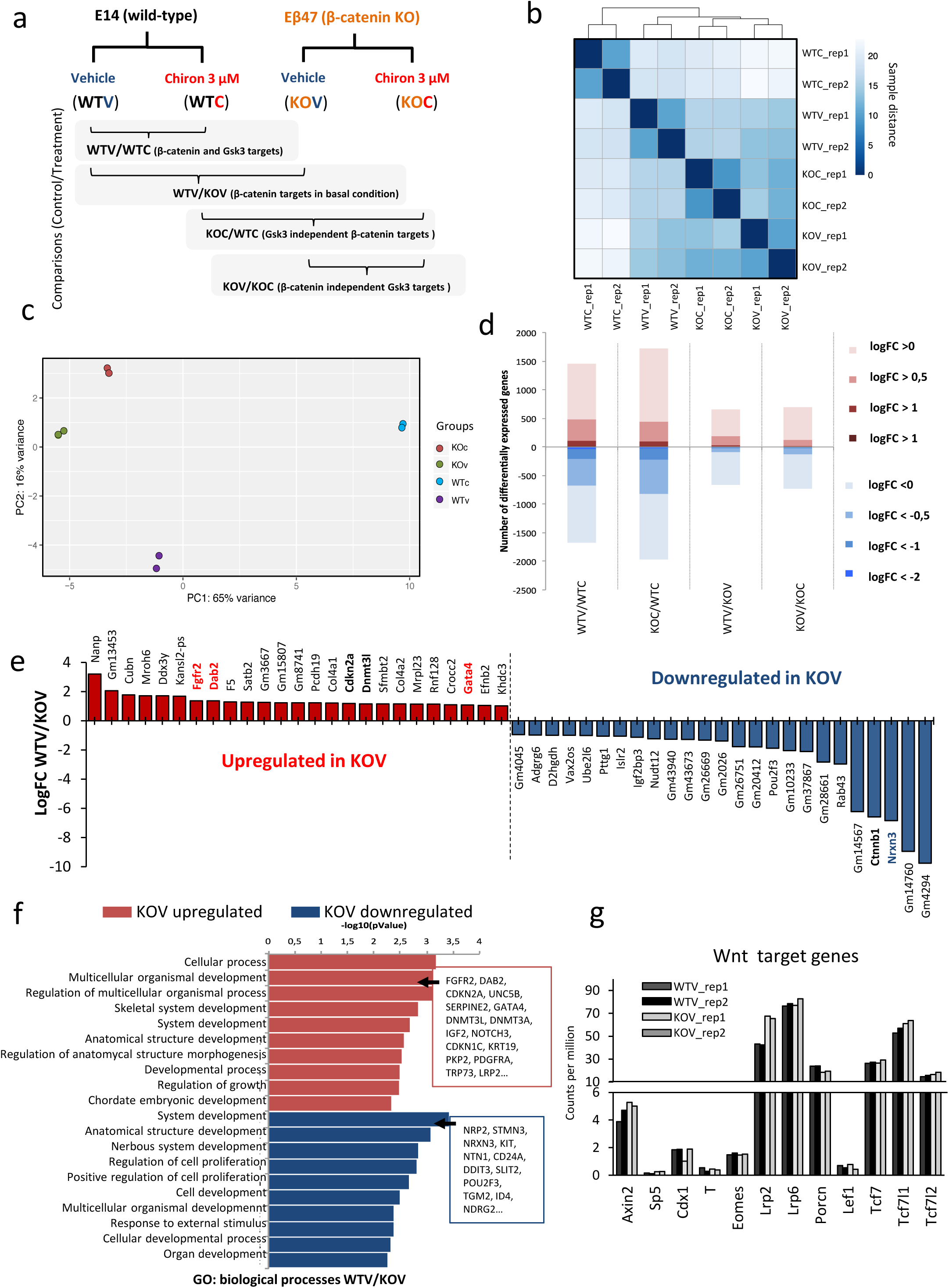
β-catenin depletion produces minor changes at transcriptomic level. **a)** Schematic representation of experimental design for RNA-seq analysis. E14 parental cells (WT) or Eβ47 cells (KO) were cultured in Serum/LIF upon 72 hours vehicle (0.3 % DMSO, V) or Chiron 3 μM (C) treatment. Two biological replicates were analysed for each sample. Pairwise sample comparisons are indicated as Control/Treatment. **b)** Sample distance matrix and hierarchical clustering of biological replicates (rep_1 and rep_2) for WTV, WTC, KOV and KOC samples. **c)** Principal component analysis plot of indicated samples. **d)** Histogram of differentially expressed genes across pairwise comparisons as indicated in Figure 3a. Shades of red indicate overexpressed genes, shades of blue indicate downregulated genes. Shade intensity represents log fold-change cut-off from >0 (no fold-change cut-off, light), to absolute log fold-change >2 (dark). Adjusted p.Value cut-off is 0.05. **e)** Top differentially expressed genes in WTV/KOV comparison ranked for log fold-change. Adjusted p-Value < 0.05. **f)** Gene ontology analysis of biological processes enriched in differentially expressed genes in WTV/KOV comparison (adjusted p.Value <0.05, absolute logFC >0.5). Upregulated features are shown in red, downregulated features are shown in blue. **g)** Histogram of RNA level (count per million reads, CPM) of canonical Wnt target genes and components across WTV (black and dark grey) or KOV (light grey and white) samples. Individual replicates are shown for each sample.

The transcriptomic profile of E14 wild-type cells cultured in Serum/LIF (WTV), was overall very similar to Eβ47, independently if cultured in presence of vehicle (KOV) or Chiron (KOC), while E14 wild-type cultured in presence of 3 μM Chiron (WTC) clustered away from all the other samples in the samples distance matrix (**Figure 3B**). Principal component analysis (PCA) showed that the highest amount of variance was due to Gsk3 inhibition in presence of β/catenin (WTC samples), while only subtle differences could be spotted between KOC, KOV and WTV samples (**Figure 3C**). These results confirmed that, similarly to what observed in β-catenin models producing N-terminally truncated isoforms [8], β-catenin absence does not significantly alter the transcriptional profile of serum/LIF cultured mESCs. We then analysed differential expressed genes (DEGs) across different samples/treatments (adjusted P. Value<0.05, logFC>0.5). Upon β-catenin depletion, 286 genes were differentially expressed with respect to E14 cells cultured in Serum/LIF+vehicle (WTV/KOV comparison, of which 192 upregulated and 94 downregulated, **Figure 3D, Figure S3B** and **Supplementary table ST1**); only 11 genes displayed a logFC higher than 2. Genes such as *Fgfr2, Dab2, Pdgfra* and *Gata4* were slightly upregulated upon β-catenin depletion although with very low fold-changes (between 1.2 and 2-fold enrichment), suggesting a partial priming toward primitive endoderm (PrE), while *Neurexin 3* (*Nrxn3*) was downregulated (**Figure 3E**). We then performed Gene ontology analysis on WTV/KOV DGEs (adjusted P. Value <0.05, logFC>0.5); both upregulated and downregulated DGEs where enriched for developmental categories such as “multicellular organism development” and “system development” (**Figure 3F, Supplementary table ST2**). Surprisingly β-catenin depletion did not alter the expression of canonical Wnt target genes such as *Axin2, Sp5, Cdx1* or *T/Brachyury* (**Figure 3G**) suggesting that canonical Wnt pathway is not active in mESCs cultured in Serum/LIF although β-catenin is highly expressed (**Figure S3A**). These results are in line with previous reports that failed to identify activity of the TOP/FOP reporter in Serum/LIF cultured mESCs [3, 4, 20] while they stand at odds with the evidence that the Wnt/β-catenin pathway is active in the inner cell mass of the blastocyst, and the canonical Wnt signalling is required for mESC self-renewal in Serum/LIF [2].

Finally, when analysing the expression levels of key lineage markers, we could not detect any change in pluripotency genes, while a slight overexpression of PrE lineage markers (*Gata4, Gata6, Foxa2, Dab2, Ihh, Cerberus* and *Pdgfra*) was observed independently of Chiron treatment in absence of β-catenin (**Figure S3C, bottom**). Such low overexpression of PrE markers upon β-catenin depletion, coupled to the absence of self-renewal and proliferation defects in serum/LIF, could either not be significant or, most likely, be due to the appearance of a rare cell subpopulation spontaneously committed toward primitive endoderm.

As colony formation and AP staining phenotypes are slightly ameliorated in β-catenin KO mESCs in presence of Chiron with respect to vehicle (**Figure 2I and S2J**), we asked whether Gsk3 inhibition *per se* could alter the transcriptome of mESCs independently of β-catenin. To address this question, we compared Eβ47 cells cultured in presence of vehicle (KOV) or 3 μM Chiron (KOC) for 72hrs (KOV/KOC comparison). Chiron treatment only resulted in minor changes as the transcriptome of KOV and KOC were overall very similar (**Figure 3B** and **3C**). Only 254 genes were differentially expressed in cells lacking β-catenin and exposed to Gsk3 chemical inhibition, again with overall low fold-change (pValue adjusted <0.05 and logFC > 0.5, of which 124 upregulated and 130 downregulated) (**Figure 3D** and **S3B, Supplementary table ST1**). Upregulated genes were enriched for biological processes such as stress-response and metabolic features, while downregulated genes were enriched for metabolic and developmental processes (**Figure S3D, Supplementary table ST3**). Nevertheless, the slight transcriptional changes induced by Chiron in absence of β-catenin revealed a certain degree of overlap with canonical Wnt target genes (**Figure S3E**). Canonical target genes such as *Myc* or *Axin2* were slightly inhibited by Chiron treatment in absence of β-catenin as well as *Tcf3* (*Tcf7l1*) mRNA levels (**Figure S3F**), the latter probably contributing to the increase in AP staining intensity observed in β-catenin KO cells exposed to Gsk3 inhibition (**Figure 2I** and **Figure S2J**). These data confirm, as previously demonstrated [21], that Gsk3 inhibition plays little or no effect on mESC transcriptional landscape independently of β-catenin. Of note Plakoglobin can partially mimic β-catenin function [22]. Although Plakoglobin protein levels are upregulated in β-catenin KO cells (**Figure 2E, S2C** and **S2D**), Plakoglobin mRNA was not found amongst the differentially expressed genes upon β-catenin loss, pointing for the existence of a post-translational regulation mechanisms for Plakoglobin which relies on β-catenin levels. In addition, our data demonstrate that Plakoglobin cannot replace nuclear β-catenin functions in response to Chiron in mESCs as no major transcriptional changes were observed in KOV/KOC comparison and, among the canonical Wnt/β-catenin targets, inhibition rather than activation was observed at best (**Figure S3E** and **S3F**).

### Canonical Wnt signalling inhibits differentiation of mESCs toward ectoderm

Although we assessed that the transcriptional effects of Chiron in absence of β-catenin are negligible, the DGEs in the KOC/WTC comparison were dependent solely on the presence of β-catenin. A similar approach has been used previously [23] by comparing mESCs treated with Chiron or XAV (a small molecule inhibitor of the Wnt pathway) without taking into account possible off-targets effects of both drugs. Our experimental approach, instead, allowed us to decouple β-catenin dependent targets (DGEs in KOC/WTC) from Gsk3-only dependent targets (DGEs in KOV/KOC).

The highest number of DEGs between comparisons represented as Control/Treatment was found in the WTV/WTC (1157 DEGs, 476 upregulated, 681 downregulated) and in the KOC/WTC (1259 DEGs, 487 upregulated, 822 downregulated) comparisons (**Figure 3D** and **S3B, Supplementary table ST1**). Wild-type mESCs treated with Chiron clustered far apart from all the other conditions, as assessed by sample distance matrix PCA (**Figure 3B** and **3C**). Numerous reports associated Wnt activation with transcriptional activation of *Nanog, Klf4, Esrrb* and *Tcfp2l1* in mESCs [17-19, 24]. However, with the exception of a slight *Tcfp2l1* upregulation and *Dppa3* downregulation, we were not able to identify any change at transcriptional levels of pluripotency marker genes upon Chiron treatment in wild-type cells (**Figure S3C**). Accordingly, we previously showed that *Nanog, Oct4* and *Rex1* levels do not change in mESCs even upon prolonged (up to 8 passages) exposure to Chiron [3]. Nevertheless, lineage markers such as *Pax6, Fgf1, Nes, Otx2, Lefty1* and *Pou3f1* were strongly inhibited by Chiron treatment in presence of β-catenin, indicating an overall reduction of differentiation commitment toward the ectoderm lineage (**Figure S3C**). Conversely, canonical Wnt target genes required for trophoectoderm and mesoderm specification, including *Cdx1, T/Brachyury*, and *Eomes*, were strongly upregulated upon Chiron treatment.

We then clustered the top 100 DGEs in the KOC/WTC comparison across the various samples (**Figure 4A**). As expected, sample clustering did not change with respect to previous analysis (**Figure 3B** and **3C**) and canonical Wnt targets such as *Axin2, Cdx1* and *T/Brachyury* were only activated in wild-type cells treated with Chiron, while little or no differences were observed in the other samples (**Figure 4A** and **4C**). Furthermore, this subset of genes was transcriptionally perturbed only in presence of both Chiron and β-catenin, once again confirming that the canonical Wnt pathway is not active in mESCs cultured in Serum/LIF unless an external stimulus (i.e. chemical Gsk3 inhibition) is applied (**Figure 4A**).

**Figure 4.**
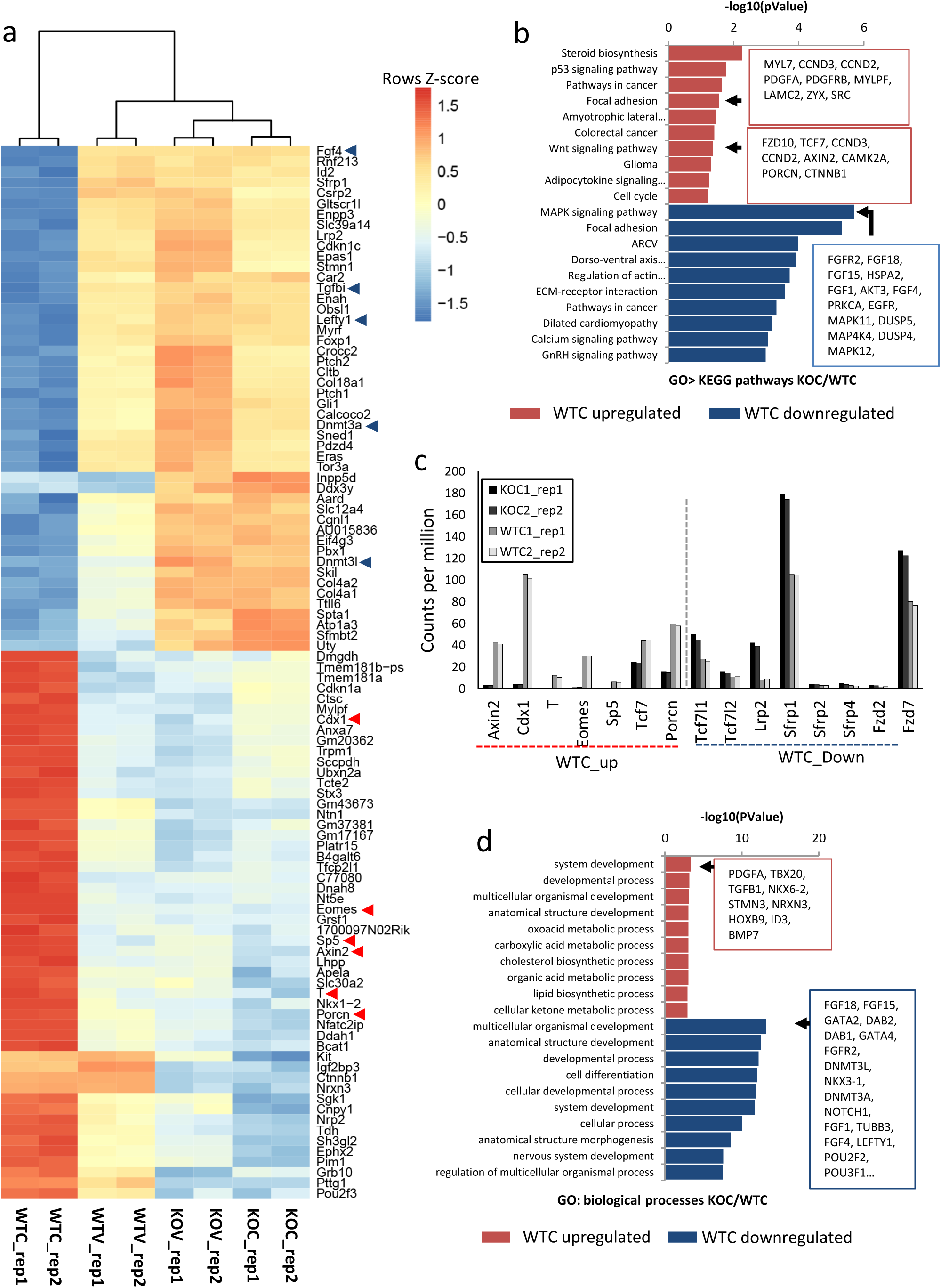
RNA-seq analysis of β-catenin dependent differentially expressed genes. Heatmap clustering of the top 100 differentially expressed genes in KOC/WTC comparison across KOV, KOC, WTV and WTC samples (p.Value adjusted <0.05, absolute logFC >0.5). Gene ontology analysis of KEGG pathways categories enriched in differentially expressed genes in the KOC/WTC comparison (p.Value adjusted <0.05, absolute logFC >0.5). **c)** Histogram of counts per million (CPMs) of canonical Wnt target genes and components differentially expressed in KOC/WTC comparison. Individual biological replicates are shown. **d)** Gene ontology analysis of biological processes categories enriched in differentially expressed genes in KOC/WTC comparison (p.Value adjusted <0.05, absolute logFC >0.5).

We next asked about the nature of transcriptional changes induced by Gsk3 inhibition in presence of β-catenin. We focused once again on the KOC/WTC comparison and performed KEGG pathways enrichment and gene ontology (DGE, s KOC/WTC, adj. pValue 0.5, logFC>0.5, **Figure 4B, 4D and Supplementary table ST5**). As expected, genes upregulated upon Chiron treatment in wild-type cells were enriched for Wnt signalling pathway category (**Figure 4B**) and known Wnt/β-catenin targets in mESCs were strongly activated by Chiron treatment in wild-type cells (**Figure 4C**). However, the Wnt signalling pathway category was also enriched in downregulated genes (**Supplementary Table ST5**), indicating an overall reassessment of all the pathways components. Interesting, Chiron treatment led to a TCF/LEFs re-assessment at nuclear level, with the downregulation of *Tcf3* (*Tcf7l1*) and the upregulation of *Tcf1* (*Tcf7*) (**Figure 4C**). Furthermore, down-regulated genes were strongly enriched for MAPK signalling pathway (**Figure 4B**) and again, to a greater extent, for focal adhesion.

Both upregulated and downregulated genes were enriched for biological processes categories associated to development. Nevertheless, the enrichment was stronger among DGEs downregulated upon Chiron treatment (**Figure 4D** and **Supplementary Table ST5**), confirming the general inhibition of spontaneous differentiation induced by the coupled action of Gsk3-inhibition and β-catenin.

### Transcriptional β-catenin activity is required to inhibit differentiation in absence of LIF

Previous reports have shown that transcriptional activity of β-catenin is dispensable for mESC self-renewal [8, 9]. In order to validate or disprove these findings, we generated lentiviruses carrying different β-catenin isoforms. In addition to the full-length β-catenin (WT βcat), a N-terminally truncated (ΔN βcat) and a C-terminally truncated isoform (ΔC βcat) were generated. While ΔN βcat isoforms mimic the isoforms generated from previous knock-out models [10], ΔC βcat carries a deletion of the transactivation domain and an impaired transcriptional activity (**Figure 5A**).

**Figure 5.**
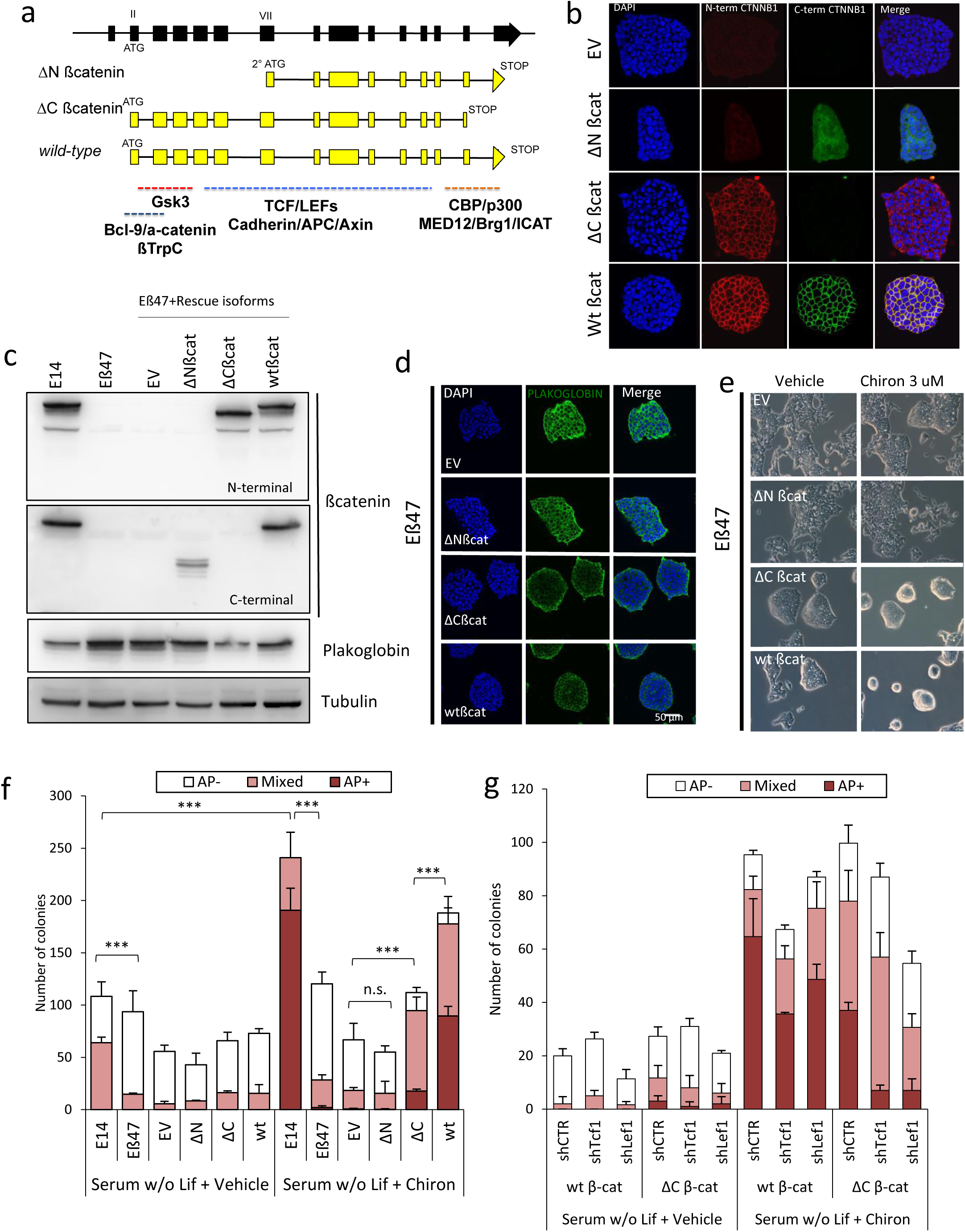
Canonical β-catenin functions are required for inhibition of differentiation. **a)** Schematic representation of β-catenin isoforms used for rescue experiments. ΔN-β-catenin mimics N-terminally truncated β-catenin isoforms obtained in previously published knock-out models. **b)** Immunofluorescence of Eβ47 cells transduced with lentiviral vectors encoding empty vector (EV), wild-type (Wt βcat) or N-terminally (ΔN βcat) and C-terminally (ΔC βcat) truncated β-catenin isoforms. Cells were stained with N-terminally (red) and C-terminally (green) β-catenin antibodies. Nuclei were counterstained with DAPI. **c)** Western blot of total protein extracts from E14 and Eβ47 un-transduced cells and Eβ47 transduced with EV, ΔN βcat, ΔC βcat and Wt βcat encoding lentiviruses. Membranes were probed with N-terminally or C-terminally raised β-catenin antibodies and anti-Plakoglobin. Tubulin was used as loading control. **d)** Immunofluorescence of Eβ47 cells transduced with EV, ΔN βcat, ΔC βcat or Wt βcat encoding lentiviruses. Cells were stained for Plakoglobin. DAPI was used to counterstain nuclei (Scalebar = 50 μm). **e)** Phase contrast pictures of Eβ47 cells transduced with EV, ΔN βcat, ΔC βcat or Wt βcat encoding lentiviruses and cultured in Serum/LIF in presence of 3 μM Chiron or Vehicle (0.3% DMSO). **f)** Alkaline phosphatase staining quantification of E14 and Eβ47 untransduced cells or Eβ47 transduced with EV, ΔN βcat, ΔC βcat or Wt βcat encoding lentiviruses. Cells were plated in Serum without LIF and supplemented with 3 μM Chiron (right) or Vehicle (0.3% DMSO, left). Error bars represent standard error of three biological replicates. Student’s t-test was used to measure statistical significance as indicated, stars indicate p-Value (n.s.=not significant, *=<0.05, **=<0.01, ***=<0.001). **g)** Alkaline phosphatase staining of Eβ47 cells transduced with either Wt βcat or ΔC βcat encoding lentiviruses. Cells were further transduced with lentivirus encoding short hairpins against Lef1 (shLef1), Tcf1 (shTcf1) or control hairpin (shCTR). Cells were cultured in Serum without LIF in presence of 3 μM Chiron or Vehicle (0.3% DMSO) for 1 week, and stained for alkaline phosphatase expression. Error bars represent standard error of three biological replicates.

Eβ47 cells were transduced with a lentivirus carrying the different β-catenin mutants under the constitutive EF1a promoter, or an empty lentivirus as a control (EV). Upon isoform expression, we studied protein subcellular localization and expression levels using N-terminal and C-terminal antibodies against β-catenin. N-terminally truncated β-catenin isoforms were only detected with the C-terminal raised antibody, and their subcellular localization recapitulated the one observed in previous knock-out models (**Figure 5B** and **Figure 1C**). Furthermore, as in ΔN-βcatenin isoforms emerging from previous knock-out models, the ΔN βcatenin overall levels were reduced compared to total β-catenin levels in wild-type cells (**Figure 5C**).

The expression and localization of ΔC βcat and WT βcat rescue isoforms were overall comparable, with intense membrane localization and similar expression levels (**Figure 5B** and **5C**). Interestingly, expression of ΔC and WT βcatenin, but not ΔN-βcatenin, restored normal Plakoglobin levels (**Figure 5C, 5D** and **S5A**), indicating that impaired membrane functions, but not the transcriptional ones, are responsible for the increased Plakoglobin levels in β-catenin knock-out cells. Similarly, ΔN βcatenin isoforms could not rescue the loss of morphological changes induced by Chiron treatment, while both ΔC and WT rescued Eβ47 cells exhibited a round shaped morphology (**Figure 5E**) indistinguishable from E14 wild-type cells upon Chiron addition (**Figure 2H**). However, only in wild-type rescued Eβ47 cells the expression of the synthetic Wnt reporter (TOP) upon Gsk3 inhibition was active (**Figure S5B**), confirming that both ΔN and ΔC β-catenin isoforms are defective for canonical Wnt signalling.

We next assessed the colony formation capacity and alkaline phosphatase expression of rescued β-catenin KO cells in different culture conditions. In Serum/LIF, only ΔC βcat and WT βcat rescued Eβ47 cells recapitulated the parental E14 cells phenotype, while ΔN βcat overexpression did not alter the phenotype with respect to empty vector transduced cells (EV, **Figure S5C, top panel**). Eβ47 showed impair clonogenicity and self-renewal, in line with previously published KO models. In serum-free media, Eβ47 could not be expanded in PD+Lif, Chiron+Lif or PD+Chiron (2i), but could self-renew in 2i/LIF; again, ΔN βcat rescued Eβ47 cells did not show any improvement of alkaline phosphatase expression or self-renewal (**Figure S5C, bottom panel**). As previously reported, both ΔC βcat and WT βcat isoforms restored self-renewal and AP staining intensity defects (**Figure S5C, bottom panel**), suggesting that canonical Wnt pathway activity is dispensable for self-renewal in these culture conditions.

However, nuclear β-catenin activity appears to be required for inhibition of differentiation in Serum culturing conditions in absence of LIF [1, 25]. Therefore, we performed clonal assay and alkaline phosphatase staining of Eβ47 rescued cells in Serum without LIF. While all the cell lines quickly differentiated in absence of LIF, Chiron treatment maintained alkaline phosphatase staining in ΔC βcat and WT βcat rescued cells, but not in ΔN βcat rescued Eβ47 (**Figure 5F** and **S5D**). Furthermore, ΔC βcat expression only partially rescued alkaline phosphatase staining, indicating that canonical Wnt activity is required to counteract spontaneous differentiation upon LIF withdrawal.

These results suggest that the inhibition of spontaneous mESC differentiation in the absence of LIF could be due to combined nuclear and membrane associated β-catenin functions. However, while ΔC β-catenin isoform fails to activate the synthetic Wnt reporter (**Figure S5B**), it has previously been observed that ΔC β-catenin isoform exhibit a certain degree of transcriptional activity, despite lacking the transactivation domain [9]. A possible explanation could be that ΔC β-catenin isoform still interact with Tcf3 and, likewise full-length β-catenin, alleviate Tcf3-mediated repression on a subset of target genes [9]. On the other hand, it has been reported that ΔC β-catenin isoforms can still form a transcriptionally active complex in association with Lef1 [26] or Tcf1. We therefore silenced Lef1 or Tcf1 in Eβ47 cells rescued with either full-length or ΔC β-catenin (**Figure S5F**) and assessed AP staining intensity upon exposure to vehicle or Chiron in absence of LIF. Tcf1 or Lef1 depletion efficiently reduced the number of AP colonies formation in response to Chiron treatment in ΔC β-catenin rescued cells and, to a lower extent, also in wild-type β-catenin mESCs (**Figure 5G** and **S5E**).

These results suggest that ΔC β-catenin could still interact with Tcf1 and Lef1, and that nuclear β-catenin/TCF/LEF functions are required to inhibit mESC spontaneous differentiation in absence of LIF, while the membrane associated β-catenin functions may have little or no role on this phenotype.

## Discussion

In this work, we showed that currently available inducible β-catenin knock-out models result in the production of N-terminally truncated isoforms in mESCs. While ΔN β-catenin isoforms are not physiologically expressed, their appearance is a consequence of rather unpredictable gene rearrangement upon genetic manipulation. By using CRISPR-Cas9 technology, we isolated three mESC clones with no detectable β-catenin expression. Of note, only a paired sgRNA approach generating a deletion encompassing the whole β-catenin gene body was successful in generating knock-out alleles (10 Kb genomic DNA deletion), while standard strategies (ORF shifting and microdeletion) only produced ΔN-truncated isoforms at best. As CRISPR-Cas9 is boosting genetic engineering, the production of undesired isoforms with potential gain-of functions, which could undermine all the conclusion of a study, should be carefully monitored and avoided. For this reason, we foresee that whole gene deletion using CRISPR-Cas9 is an advantageous approach for generating knock-out models free of undesired side products.

We assessed that the appearance of ΔN-truncated β-catenin isoforms in previously published β-catenin KO mESC models is not detrimental for their phenotype assessment, as no compensatory or overlapping effects were to be attributed to the truncated proteins. In addition, none of the impaired phenotypes observed upon complete β-catenin loss could be rescued by a ΔN-β-catenin isoform, thus recapitulating the phenotype observed in the previously generated KO models.

β-catenin depletion does not alter self-renewal and pluripotency marker expression of mESCs under serum/LIF culturing condition, with expected morphological changes probably rescued by Plakoglobin stabilization. Moreover, β-catenin depletion also does not alter the expression of known canonical Wnt targets in basal culturing conditions suggesting that the Wnt/β-catenin pathway is not constitutively active in mESCs cultured in feeder-free media supplemented with Serum/LIF. Accordingly, β-catenin loss does not globally alter the transcriptional profile of mESCs but induces a mild activation of primitive endoderm marker genes, which could potentially highlight the presence of a rare population of terminally differentiated cells that cannot withstand β-catenin absence.

Furthermore, only chemical inhibition of Gsk3 activates a transcriptional Wnt response in wild-type cells, while eliciting little or no response on β-catenin knock-out mESCs, suggesting that Plakoglobin upregulation cannot recapitulate nuclear β-catenin functions. Nevertheless, activation of the canonical Wnt pathway shields mESCs from spontaneous epiblast/ectoderm differentiation by suppressing the expression of lineage specific genes and not by enhancing the expression levels of pluripotency factors. This effect is particularly evident when cells are deprived of LIF, suggesting overlapping functions between LIF and Wnt signalling in mESC self-renewal. With respect to these functions, Chiron treatment can prolong, but not indefinitely sustain, mESC self-renewal in absence of LIF. This phenotype is abrogated by β-catenin loss but can be rescued by wild-type β-catenin and, to a lower extent, by ΔC-β-catenin isoforms, posing the question of whether nuclear β-catenin functions are truly required for differentiation inhibition. We demonstrated that Tcf1 and Lef1 depletion further impair the colony formation capacity and alkaline phosphatase staining of both wild-type and ΔC rescued β-catenin KO cells, suggesting that ΔC-β-catenin isoforms are not completely transcriptionally silent and can still interact not only with Tcf3 (as previously suggested [9]) but also with other TCF/LEFs. Finally, we proved that activation of the canonical Wnt pathway sustains mESC self-renewal through inhibition of spontaneous differentiation and that nuclear β-catenin functions, in associations with TCF/LEFs family members, are required for this phenotype.

In the future, it will be particularly interesting to investigate further the transcriptional response of ΔC-rescued β-catenin KO cells to Chiron, as it could pinpoint a subset of powerful transcriptional targets truly responsible for this Wnt-mediated self-renewal enhancement.

## Materials and methods

### Cell culture

Mouse ESCs E14 (129/Ola strain) were cultured on gelatin-coated plates in ESC medium: DMEM supplemented with 15% FBS (Hyclone), 1X non-essential amino acids, 1X GlutaMax, 1X penicillin/ streptomycin, 1X 2-mercaptoethanol and 1,000 U/ml LIF ESGRO (Chemicon). mESCs cultured in Serum+Lif medium were replated evey 3 days at a split ratio from 1:30 following dissociation with Trypsin 0.05% EDTA (Gibco). mESCs containing floxed alleles of b-catenin were a kind gift of professor Rolf Kemler (SR18 and NLC1 cell lines, stably expressing CRE-ERT2) and professor Christine Hartmann (Ctnnb1^fl/fl^ and Ctnnb1^del/del^).

For serum-free cultures, mESCs were cultured without feeders or serum in pre-formulated N2B27 medium (NDiff™ N2B27 base medium, Stem Cell Sciences Ltd, Cat. No. SCS-SF-NB-02) supplemented with Small molecule inhibitors PD0325901 (PD, 1 μM, Selleck) and CHIRON99021 (CH, 3 μM, Selleck) and 1000 U/ml LIF (ESGRO, Millipore). Cells were routinely propagated on 0.1% gelatin-coated plastic and replated every 3 days at a split ratio of 1 in 10 following dissociation with Accutase (Gibco) as previously reported [9].

Human embryonic kidney 293t (HEK293t) were purchased from ATCC (293T (ATCC^®^ CRL-3216™) and cultured in DMEM supplemented with 10% FBS (Hyclone), 1X penicillin/ streptomycin. HEK293t were replated every 3 days at a split ratio of 1 in 6 following dissociation with Trypsin 0.05% EDTA (Gibco, Life technologies).

### qRT-PCR

RNA was extracted and purified using with Maxwell LEV semi-automated RNA extraction kit (Promega) following manufacturer instructions. The cDNA was produced with iScript cDNA synthesis kit (BioRad). Real-time quantitative PCR reactions from 8,3 ng of cDNA were set up in triplicate using a LightCycler DNA SYBR Green I Master PCR machine (Roche). The oligonucleotides used in qRT-PCR experiments are provided in **SI6**.

### Western blot, IF and flow cytometry staining

For western blot experiments cells were harvested and washed twice with PBS. Cell lysis was performed on ice for 25 min, in RIPA buffer (150 mM NaCl, 1% Nonidet P40, 0.5% sodium deoxycholate, 0.1% sodium dodecyl sulphate, 50 mM Tris-HCl, pH 8.0) containing a protease inhibitory cocktail (Roche). Insoluble material was pelleted by centrifugation at 16,000× g for 3 min at 4 °C. Protein concentrations were determined using the Bradford assay (Bio-Rad). Thirty micrograms extract was mixed with 4× sample buffer (40% glycerol, 240 mM Tris/HCl, pH 6.8, 8% SDS, 0.04% bromophenol blue, 5% β-mercaptoethanol), denatured at 96°C for 5 minutes, separated by SDS-PAGE, and transferred to nitrocellulose membranes (PROTRAN-Whatman, Schleicher&Schuell). The membranes were blocked with 5% non-fat dry milk in TBS-T for 60 min, incubated with primary antibodies overnight at 4 °C, washed three times with TBS-T for 10 min, incubated with the peroxidase-conjugated secondary antibody (1:2000; Amersham Biosciences) in TBST with 5% non-fat dry milk for 60 min, and washed three times with TBST for 10 min. Immunoreactive proteins were detected using Supersignal West Dura HRP Detection kits (Pierce). A list of the primary antibodies is provided in **SI5**.

For cytometry analysis, cells were trypsinised, washed once in PBS, resuspended in PBS with 5% FBS + DAPI and analysed on BD Fortesa cytometer. For E-cadherin staining incubation with the PE conjugated antibody was performed after the first wash. Neuronally differentiated mESCs were used as negative staining control.

For immunocytochemistry, mESCs were fixed with 4% paraformaldehyde for 20 min at room temperature, and then washed twice with PBS following incubation in blocking solution containing 10% goat serum or 3% Bovine Serum Albumin (Sigma) and 0.1% Triton X-100 (Sigma) for 1 h at room temperature. The cells were then left overnight at 4 °C in blocking solution containing the primary antibody. The next day, the cells were washed three times with PBS and then incubated with the secondary antibody for 1 h at room temperature in PBS. The primary antibodies used are provided in **SI5**. Goat anti-mouse IgG, goat anti-rabbit IgG, (1:1000, Life Technologies) conjugated to Alexa Fluor-488 or Alexa Fluor-594 were used as secondary antibodies. Nuclear staining was performed with DAPI (Life Technologies).

### RNA-seq

Total RNA from mESCs was extracted wit RNAeasy kits (QIAGEN) following manufacturer’s instructions. RNA integrity check, Poly-A pull-down and library preparation were performed by CRG genomic facility. Samples were sequenced in two biological replicates to a depth of 30 million reads (100 bp) per sample using Illumina HiSeq Reads were mapped to a reference transcriptome (mouse transcriptome from Ensembl v80) using kallisto (v0.42.5) [27], to generate counts and “transcripts per million” (TPM) values. We tested for differentially expressed genes for all pairwise conditions using edgeR [28] from raw counts, filtering for genes with p values < 0.001 and a fold change greater than 1. GO and KEGG analysis were performed using DAVID (6.8 beta version) [29, 30]. Raw data are available upon request.

### Constructs

sgRNAs were cloned by annealed oligos cloning into px459-SpCas9-Puro (Addgene # 48138) as previously described [3], a list of the oligos used for generating Ctnnb1 targeting vectors is listed in **SI3**. pSpCas9(BB)-2A-GFP (PX458) was a gift from Dr. Feng Zhang. Lentiviral vectors expressing wt or truncated β-catenin isoforms (pL-EF1a-Puro) were generated by Gibson assembly. pL-EF1a-Puro backbone by PCR on 7TGP vector (Addgene#24305) while hEF1a promoter was amplified from p1494 EF1a Ires Hygro vector [4]. Wt β-catenin CDS was amplified from mESCs cDNA using Superscript III first-strand cDNA synthesis kit. Mutant β-catenin CDSs were generated by PCR on the wild-type product. Sequences for pL-EF1a Puro based vectors expressing wt and mutant β-catenin isoforms are provided in **SI2**. 7TGP was a gift from Dr. Roel Nusse. Short-hairpins RNAs were cloned in pLKO Hygro as previously described [4]. Oligonucleotides sequences for short-hairpin cloning are provided in **SI4**. pLKO.1 hygro was a gift from Bob Weinberg (Addgene plasmid # 24150). The maps and the full-length sequences of all the constructs generated in this study are provided in **SI2**.

### Generation of Ctnnb1 KO cells

CRISPR/Cas9 was used to induce small in-dels, microdeletions or complete deletion of the Ctnnb1 locus using single sgRNA or pairwise combinations. Briefly for each experiment 5×10^^6^ mESCs (E14Tg2a from ATCC) per well were seeded onto gelatin-coated mw6 plates. 24 hours after seeding, 2 ml of fresh mESCs medium were provided at least 30 minutes before transfection. Transfection mix consisted of 5 ug of all-in-one vectors expressing Cas9 and previously subcloned sgRNA (px459-spCas9-Puro), 100 ul Optimem (Thermo-Fisher) and 20 ul Polyfectamine reagent (Qiagen). For co-transfection of two sgRNAs, 2.5 ug of each vector were used. Transfection mix was incubated 15 minutes at room temperature and then directly added to seeded mESCs. Fresh mESCs medium was added to a final volume of 2,7 ml and 24 hours after, medium was replaced. 48 hours after transfection Puromyicin selection (5 ug/ml) was applied for additional 48 hours. Cells were then analysed at population level to assess the knock-out efficiency. For establishment of mESCs ß-catenin KO cell lines, transfected pools were replated at clonal density and single cell clones were manually picked and screened for homozygous Ctnnb1 deletion. For PCR assay of Ctnnb1 two or three oligonucleotides were used depending on the deletion strategy. Knock-out and screening strategies together with oligos for genotyping edited cells are summarised in **SI4**.

### Cell cycle analysis

For cell cycle analysis of mESCs, cells were detached with Accutase (Gibco) and collected by centrifugation at 300 rcf for 5 min. The cell pellet was resuspended and fixed overnight in 3ml ice-cold 70% ethanol. After fixation, the cells were centrifuged at 300rcf for 10 min at room temperature. The pellet was washed twice in 1ml PBS. During each wash, the cells were pelleted at 300rcf for 5 min at room temperature. Then cells were resuspended in DAPI solution (SIGMA 09542) (5μg/ml/106 cells in PBS) and incubated for 30 min on ice. Samples were processed with BD LSR Fortessa and analysed with FlowJo software.

### Lentivirus production

For mESCs transduction, lentiviral particles were produced following the RNA interference Consortium (TRC) instructions for lentiviral particle production and infection in 6-well plates (http://www.broadinstitute.org/rnai/public/). Briefly, 5 ×105 HEK293T cells/well were seeded in 6-well plates. The day after plating, the cells were co-transfected with 1 μg of lentiviral vector, 750 μg pCMV-dR8.9, and 250 μg pCMV-VSV-G, using Polyfect reagent (Qiagen). The day after transfection, the HEK293T culture medium was substituted with the ESC culture medium. Then 5 ×105 ESCs/well were plated onto gelatin-coated 6-well plates the day before transduction. The lentiviral-particles containing medium was harvested from HEK293T cells at 48, 72 and 96 h after transfection, filtered, and added to the ESC plates. The day after transduction, these ESCs were washed twice in PBS and hygromycin selection (50 μg/ml) was applied.

## Figure legends

**Figure S1.**
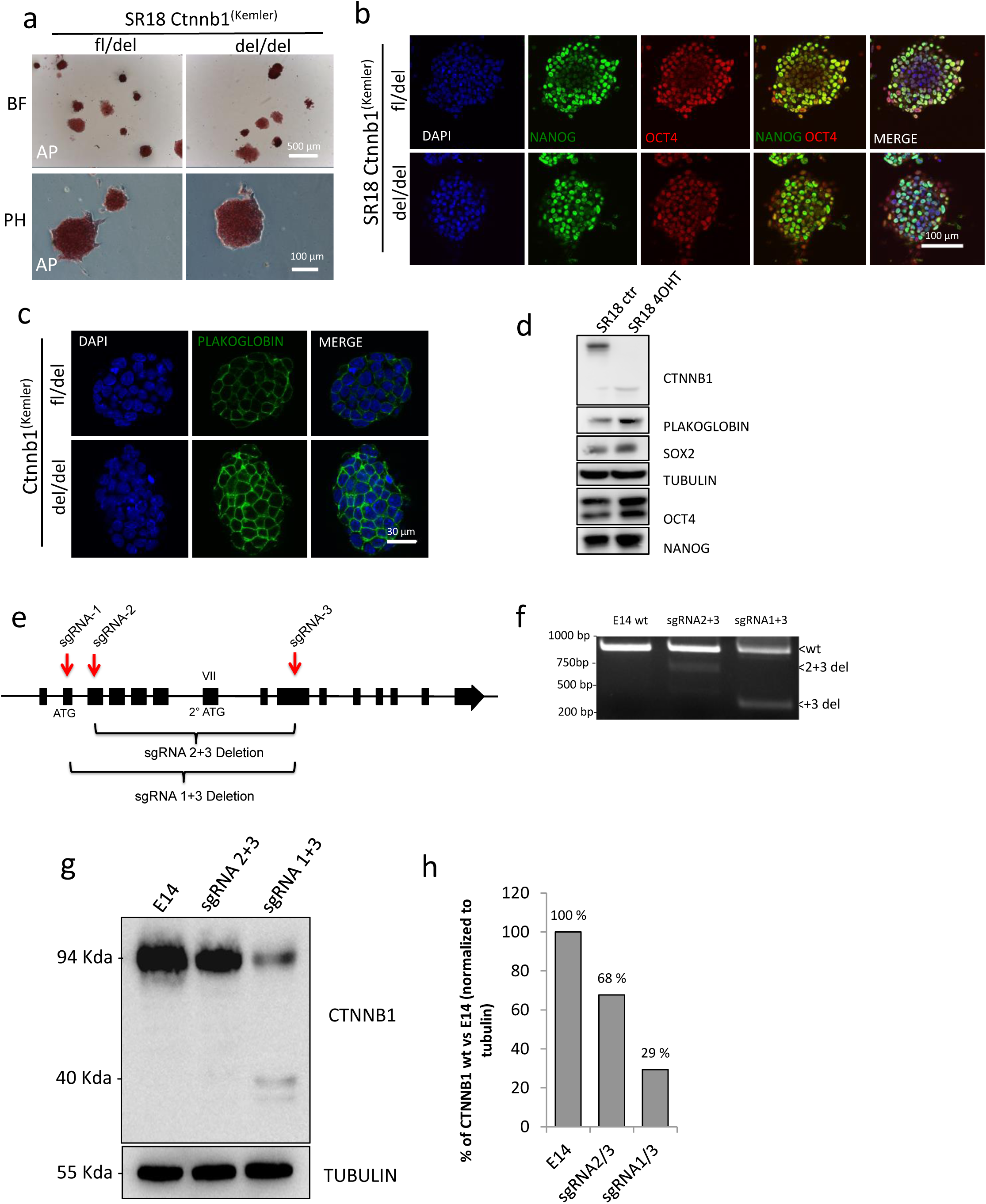
Knock-out models producing N-terminally truncated isoforms display normal clonogenicity and pluripotency marker expression. **a)** SR18 Ctnnb1fl/del or Ctnnb1 del/del display overall similar alkaline phosphatase staining expression and morphology. BF=brightfield (scalebar=500 μm), PH=phase contrast (scalebar=100 μm). **b)** Immunofluorescence of Nanog and Oct4 on fixed SR18 Ctnnb1^fl/del^ or Ctnnb1^del/del^ cells. Scalebar=100 μM. DAPI was used to counterstain nuclei. **c)** Immunofluorescence of Plakoglobin on fixed SR18 Ctnnb1^fl/del^ or Ctnnb1^del/del^ cells. Scalebar=30 μM. DAPI was used to counterstain nuclei. **d)** Western blot of total protein extracts of SR18 Ctnnb1fl/del (SR18 ctr) or Ctnnb1 del/del (SR18 4OHT) cells. Protein extracts were probed for β-catenin (CTNNB1), Plakoglobin, Sox2, Oct4 and Nanog. Tubulin was used as loading control. **e)** Schematic representation of sgRNAs target positions along the β-catenin locus. sgRNAs were used in pairwise combinations to excise different gene regions. sgRNAs are represented as red arrows, indicating the position and orientation of oligonucleotides used for PCR genotyping (3 oligos PCR). **f)** PCR-genotyping of E14 mESCs transiently transfected with Cas9 and pairwise combinations of sgRNAs as depicted in e). Untransfected cells were used as parental control. Expected amplicon size is 824 bp for wild-type, 595 bp for sgRNA2+sgRNA3, 278 bp for sgRNA1+sgRNA3. **g)** Western blot for β-catenin on total protein extract of E14 mESCs parental cell line, or upon transient transfection of Cas9 and pairwise combination of sgRNAs as in e). Tubulin was used as a loading control. **h)** Quantification of full-lenght β-catenin deletion in g). β-catenin band intensity was normalized on Tubulin intensity for each sample and then rescaled as a percentage of the untrasfected parental cell line.

**Figure S2.**
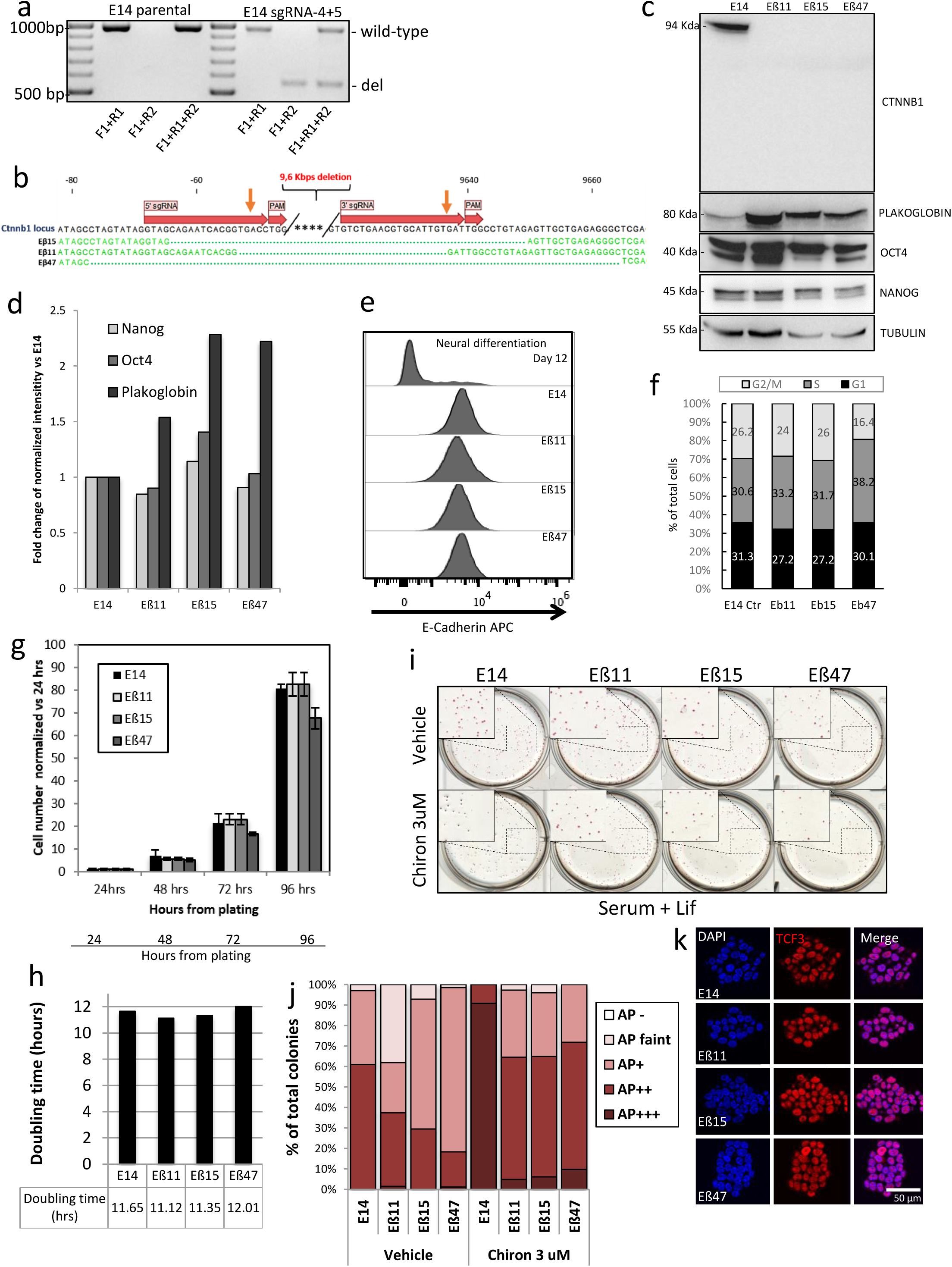
Characterization of β-catenin knock-out clones. **a)** PCR genotyping of E14 transiently transfected with Cas9, sgRNA4 and sgRNA5 (right) and parental cell line (left). Three different oligos combinations were used to detect wild-type allele (F1+F2), deleted alleles (F1+R2) or both (F1+R1+R2). **b)** Sanger sequencing of Eβ11, Eβ15 and Eβ47 Ctnnb1 edited locus. Matching bases are represented as green letters, green dots are deleted bases. Orange arrows indicate expected Cas9 editing sites, sgRNAs sequences and PAM are shown as red arrows. **c)** Western blot of total protein extracts from E14, Eβ11, Eβ15 and Eβ47 mESCs. Protein extracts were probed for β-catenin, Plakoglobin, Nanog and Oct4 expression. Tubulin was used as loading control. **d)** Band intensity quantification relative to western-blot in figure (c). Band intensities were normalized on Tubulin intensity for each sample and then rescaled as fold-change with respect to the parental cell line. **e)** Flow cytometry analysis of E-Cadherin expression in Eβ11, Eβ15, Eβ47 and parental E14 cells. E14 cells undergoing neuro-ectodermal differentiation were used as negative control for E-Cadherin expression (top). **f)** Flow cytometry cell-cycle analysis on fixed E14, Eβ11, Eβ15 and Eβ47 cells. DAPI was used to measure DNA content. Data are represented as histogram depicting the percentage of cells in G1 (black), S (grey) or G2/M (light grey). One exemplificative experiment. **g, h)** Growth curve of E14, Eβ11, Eβ15 and Eβ47 cells and doubling time analysis. **i)** AP staining of E14, Eβ11, Eβ15 and Eβ47 cells cultured in Serum/LIF in presence of Vehicle (0.3 % DMSO) or 3 μM Chiron for 5 days. Whole plate scanning and magnification inset (dashed boxes). **j)** AP staining intensity quantification relative to Figure S2i. **k)** Immunofluorescence of parental E14 cells, Eβ11, Eβ15 and Eβ47 for Tcf3 expression. DAPI was used to counterstain nuclei. Scalebar= 50 μm.

**Figure S3.**
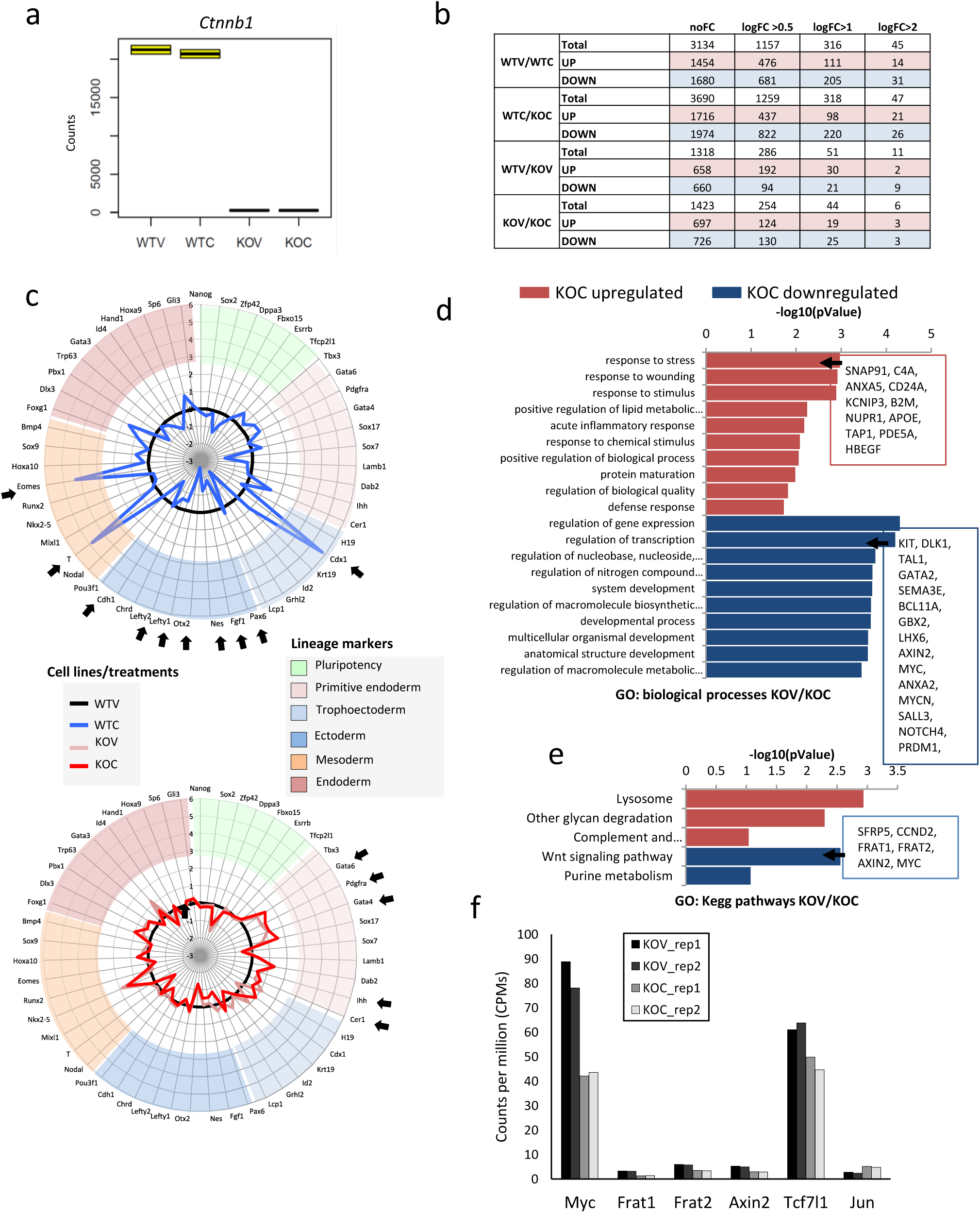
RNA-seq analysis of β-catenin depletion and Gsk3 inhibition in mESCs. **a)** Box plot of Ctnnb1 mRNA expression levels (raw counts) across WTV, WTC, KOV and KOC samples. **b)** Number of differentially expressed genes in various comparison relative to Figure 3d. **c)** Radar plot showing the fold-change of pluripotency and lineage marker genes in WTC (top panel, blue line) or KOV, KOC samples (light and dark red lines respectively, bottom panel), versus WTV sample (black line, top and bottom panel). **d, e)** Gene ontology analysis of biological processes (d) and KEGG pathways (e) enriched in differentially expressed genes in the KOV/KOC comparison (adjusted p.Value <0.05, absolute logFC >0.5) **f)** Histogram of counts per million (CPMs) of canonical Wnt target genes with minor expression level changes in KOV/KOC comparison. Individual biological replicates are shown.

**Figure S5.**
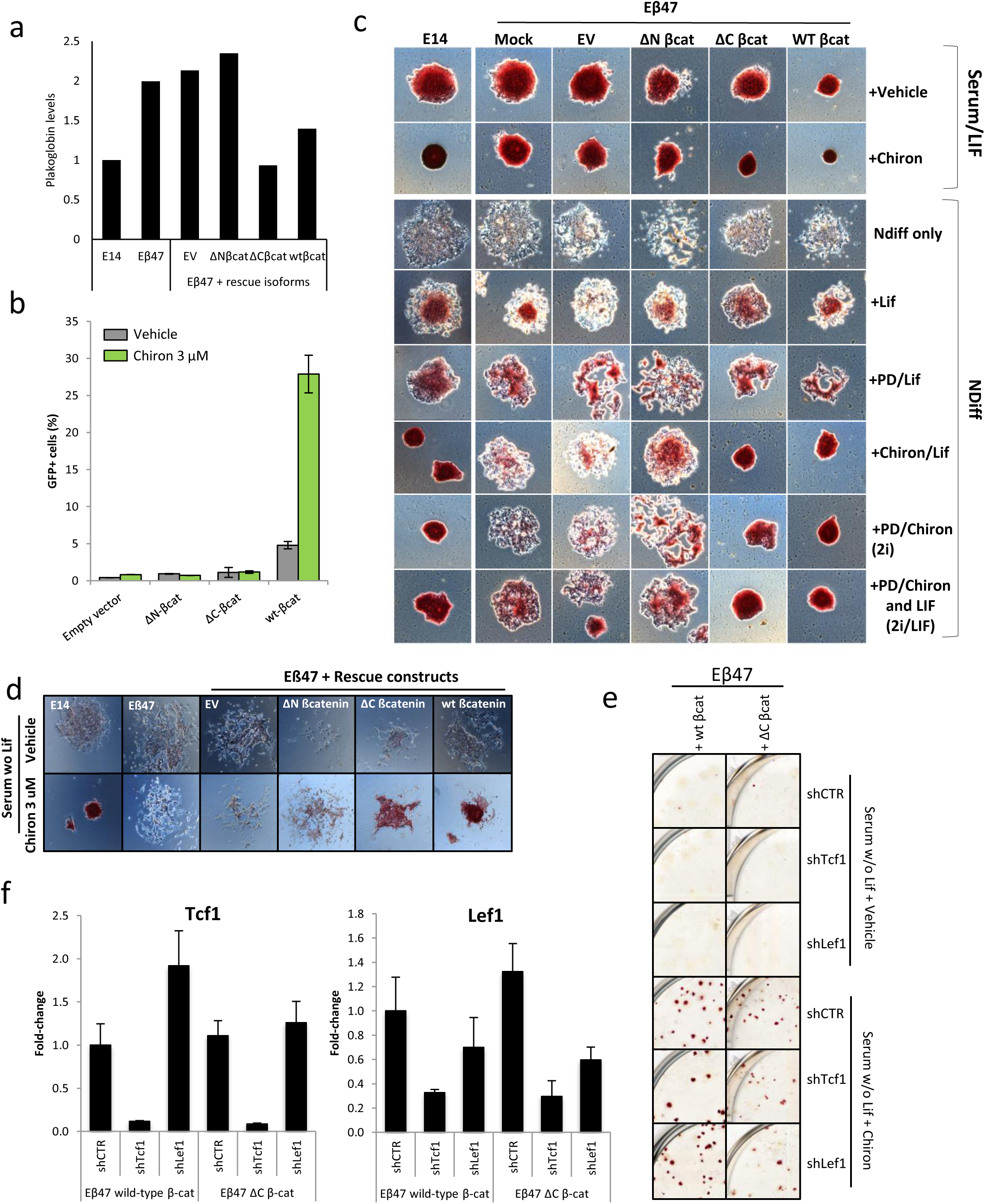
Canonical β-catenin functions are required for inhibition of differentiation. **a)** Western blot quantification of Plakoglobin levels relative to Figure 5c. Plakoglobin/Tubulin ratios are represented as fold change with respect to the E14 sample. **b)** Eβ47 cells were transduced with either EV, ΔN βcat, ΔC βcat or Wt βcat encoding lentiviruses. Cells were further transduced with the 7TGP Wnt reporter lentivirus and cultured for 48 hours in presence of 3 μM Chiron (green bars) or Vehicle (0.3 % DMSO, grey bars). The percentage of eGFP positive cells is represented for each sample. Error bars represents standard error of two technical replicates. **c)** Alkaline phosphatase staining of E14 and Eβ47 cells, and Eβ47 cells, transduced with rescue β-catenin isoforms encoding lentiviruses and cultured for 1 week in the indicated media. **d)** Exemplificative phase contrast pictures relative to Figure 5f. **e)** Alkaline phosphatase staining exemplificative pictures relative to Figure 5g. **f)** qRT-PCR of Tcf1 and Lef1 levels in Eβ47 cells rescued with either Wt βcat or ΔC βcat plasmids and transduced with short hairpin encoding lentiviruses against Tcf1 or Lef1. Error bars represents standard errors of technical duplicates.

## Author Contribution

FA, LM and MPC wrote the manuscript; FA and MP designed the experiments. MPC supervised the project. FA, FS and EP performed the experiments.

## Acknowledgments

This work was supported by European Union’s Horizon 2020 Research and Innovation Programme [CellViewer No 686637 to M.P.C.]; the Ministerio de Economia y Competitividad y FEDER (SAF2011-28580, and BFU2015-71984 to M.P.C.), an AGAUR grant from Secretaria d’Universitats i Investigació del Departament d’Economia i Coneixement de la Generalitat de Catalunya (2014SGR1137 to M.P.C.). The Spanish Ministry of Economy and Competitiveness (MEIC), the EMBL partnership, Centro de Excelencia Severo Ochoa 2013-2017; the CERCA Programme/Generalitat de Catalunya (M.P.C) and Ministerio de Ciencia e Innovación FPI (to F.A.), La Caixa international PhD fellowship (to F.S.); KU Leuven C1 funds (C14/16/078) and FWO (G097618N) funds to F.LL. The funders had no role in study design, data collection and analysis, decision to publish, or preparation of the manuscript.

