## Supplementary information for "Canonical Wnt pathway controls mESCs self-renewal through inhibition of spontaneous differentiation via β-catenin/TCF/LEF functions"

**SI1 - Sanger sequencing**
Results relative to **Figure S2 B**
Forward sequencing oligo (5’-3’): CACCGTATGCCTACAATCTGTTTCTA
Reverse sequencing oligo (5’-3’): CTACACAATGTTACACGTCTCCAGAT

**>Eβ11-Forward**
NNNNNNNNCTTCCTGACCCGTGGCTGCTGTGTATTTTTAGTGTATGCCATGGTGAACTGGCTTTTGGTGTCGGGGCA
CATAGCCTAGTATAGGTAGAGTTGCTGAGAGGGCTCGAGGGGTGGGCTGGTATCTCAGAAAGTGCCTGACACACTAA
CCAAGCTGAGTTTCCTATGGGAACAGTCGAAGTACGCTTTTTGTTCTGGTCCTTTTTGGTCGAGGAGTAACAATACAAA
TGGATTTGGGGAGTGACTCACGCAGTGAAGAATGCACACGAATGGATCACAAGATGGCGTTATCAAACCCTAGCCTT
GCTTGTTCTTTGTTTTAATATCTGTAGTGGTGCTGACTTTGCTTGCTTTTATTTTTTGCAGTAACTGTTAGTTTTTAAGTA
GTGTTATGTTCTAGTGAACCTGCTACAGCAATTTCTGATTTCTAAGAACCGAGTAATGGTGTAGAACACTAATTCATAA
TCACGCTAATTGTAATCTGGAGACGTGTACAATTGTGTAGANNNNA

**>Eβ11-Reverse**
TATAGCGTGATTTGATTCGTGTTCTACACCATTTACTCCGGTTTCTTAGAAATCAGAAATTTGCTGTAGCAGGTTCACTA
GAACATAACACTACTTAAAAACTAACAGTTACTGCAAAAAATAAAAGCAAGCAAAGTCAGCACCACTACAGATATTAA
AACAAAGAACAAGCAAGGCTAGGGTTTGATAACGCCATCTTGTGATCCATTCGTGTGCATTCTTCACTGCGTGAGTCA
CTCCCCAAATCCATTTGTATTGTTACTCCTCGACCAAAAAGGACCAGAACAAAAAGCGTACTTCGACTGTTCCCATAGG
AAACTCAGCTTGGTTAGTGTGTCAGGCACTTTCTGAGATACCAGCCCACCCCTCGAGCCCTCTCAGCAACTCTACCTAT
ACTAGGCTATGTGCCCCGACACCAAAAGCCAGTTCACCATGGCATACACTAAAAATACACAGCAGCCACGGTGTCAGG
AAGCTCTTCTCAGTAGAAACAGATTGTAGGCATAACGGTGA

**>Eβ15 Forward**
CCCGNNANNNNNNNNGNCNNGTGGCTTGCNGTGTATTTTNAGTTGTATGCCATGGTGAACTGGCTTTTGGTGTCGG
GGCACATAGCCTAGTATAGGTAGCAGAATCACGGGATTGGCCTGTAGAGTTGCTGAGAGGGCTCGAGGGGTGGGCT
GGTATCTCAGAAAGTGCCTGACACACTAACCAAGCTGAGTTTCCTATGGGAACAGTCGAAGTACGCTTTTTGTTCTGG
TCCTTTTTGGTCGAGGAGTAACAATACAAATGGATTTGGGGAGTGACTCACGCAGTGAAGAATGCACACGAATGGAT
CACAAGATGGCGTTATCAAACCCTAGCCTTGCTTGTTCTTTGTTTTAATATCTGTAGTGGTGCTGACTTTGCTTGCTTTT
ATTTTTTGCAGTAACTGTTAGTTTTTAAGTAGTGTTATGTTCTAGTGAACCTGCTACAGCAATTTCTGATTTCTAAGAAC
CGAGTAATGGTGTAGAACACTAATTCATAATCACGCTAATTGTAATCTGGAGACGTGTACAATTTGTGTAGANNNNA

**>Eβ15-Reverse**
NNNNNNNNNGTGNNNTGANNNGTNGTNCTACACCATTTACTCCGGTTNCTTAGAAATCAGAAATTNGCTGTAGCAG
GTTCACTAGAACATAACACTACTTAAAAACTAACAGTTACTGCAAAAAATAAAAGCAAGCAAAGTCAGCACCACTACA
GATATTAAAACAAAGAACAAGCAAGGCTAGGGTTTGATAACGCCATCTTGTGATCCATTCGTGTGCATTCTTCACTGC
GTGAGTCACTCCCCAAATCCATTTGTATTGTTACTCCTCGACCAAAAAGGACCAGAACAAAAAGCGTACTTCGACTGTT
CCCATAGGAAACTCAGCTTGGTTAGTGTGTCAGGCACTTTCTGAGATACCAGCCCACCCCTCGAGCCCTCTCAGCAACT
CTACAGGCCAATCCCGTGATTCTGCTACCTATACTAGGCTATGTGCCCCGACACCAAAAGCCAGTTCACCATGGCATAC
ACTAAAAATACACAGCAGCCACGGTGTCAGGAAGCTCTTCTCAGTAGAAACAGATTGTAGCCNTTACGGGTGANNA

**>Eβ47-Forward**
NNNGNNNNNCTNCCTGACCNGTGGCTGCTGTGTATTTTTAGTGTATGCCATGGTGAACTGGCTTTTGGTGTCGGGGC
ACATAGCTCGAGGGGTGGGCTGGTATCTCAGAAAGTGCCTGACACACTAACCAAGCTGAGTTTCCTATGGGAACAGT
CGAAGTACGCTTTTTGTTCTGGTCCTTTTTGGTCGAGGAGTAACAATACAAATGGATTTGGGGAGTGACTCACGCAGT
GAAGAATGCACACGAATGGATCACAAGATGGCGTTATCAAACCCTAGCCTTGCTTGTTCTTTGTTTTAATATCTGTAGT
GGTGCTGACTTTGCTTGCTTTTATTTTTTGCAGTAACTGTTAGTTTTTAAGTAGTGTTATGTTCTAGTGAACCTGCTACA
GCAATTTCTGATTTCTAAGAACCGAGTAATGGTGTAGAACACTAATTCATAATCACGCTAATTGTAATCTGGAGACGTG
TACATTNGTGTAGANNNA

**>Eβ47-Reverse**
NNTNNNNCGTGATTATGATTAGTGTTCTACACCATTACTCGGTTCTTAGAAATCAGAAATTGCTGTAGCAGGTTCACT
AGAACATAACACTACTTAAAAACTAACAGTTACTGCAAAAAATAAAAGCAAGCAAAGTCAGCACCACTACAGATATTA
AAACAAAGAACAAGCAAGGCTAGGGTTTGATAACGCCATCTTGTGATCCATTCGTGTGCATTCTTCACTGCGTGAGTC
ACTCCCCAAATCCATTTGTATTGTTACTCCTCGACCAAAAAGGACCAGAACAAAAAGCGTACTTCGACTGTTCCCATAG
GAAACTCAGCTTGGTTAGTGTGTCAGGCACTTTCTGAGATACCAGCCCACCCCTCGAGCTATGTGCCCCGACACCAAA
AGCCAGTTCACCATGGCATACACTAAAAATACACAGCAGCCACGGTGTCAGGAAGCTCTTCTCAGTAGAAACAGATTG
TAGCCNTNACCGGTGANNA

#### SI2 - Constructs maps and sequences

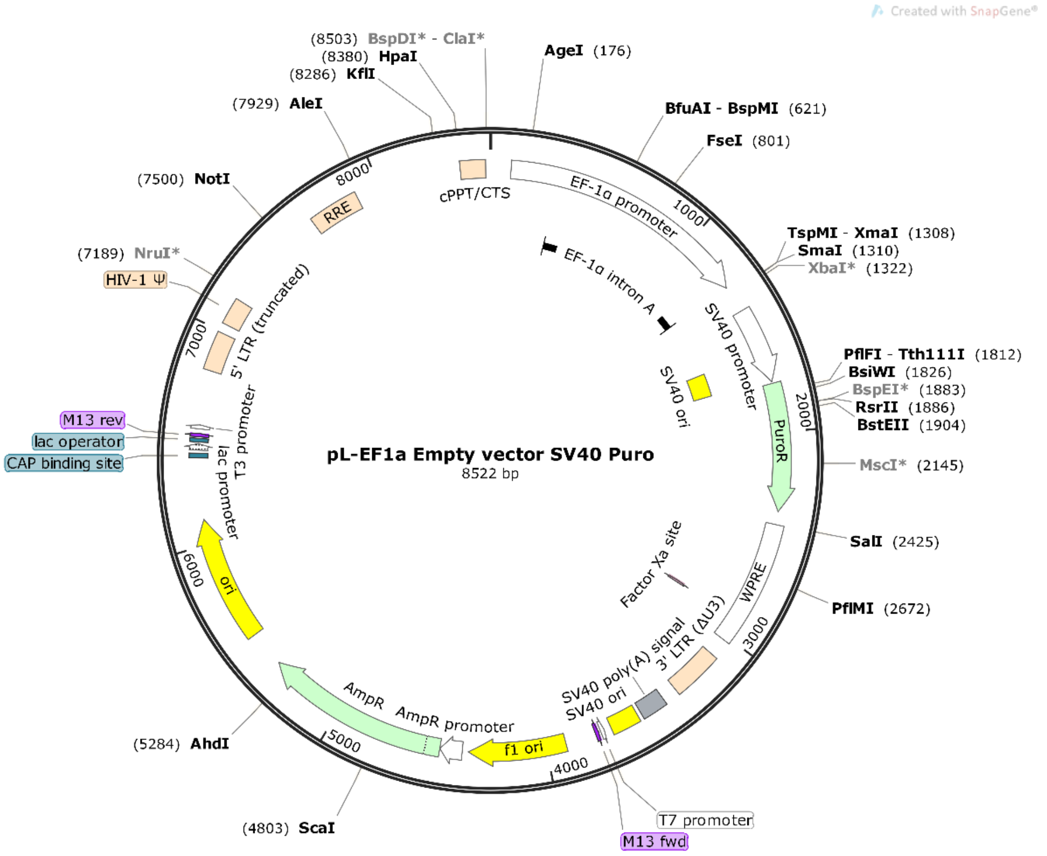

#### >pL-EF1a empty vector SV40 Puro

CTAGCCCCGATAAGCTTTGCAAAGATGGATAAAGTTTTAAACAGAGAGGAATCTTTGCAGCTAATGGACCTTCTAGGTCTTGAAAGGAGTGGGAATTGGCTCCGGTGCCCGTCA
GTGGGCAGAGCGCACATCGCCCACAGTCCCCGAGAAGTTGGGGGGAGGGGTCGGCAATTGAACCGGTGCCTAGAGAAGGTGGCGCGGGGTAAACTGGGAAAGTGATGTCGTGTA
CTGGCTCCGCCTTTTTCCCGAGGGTGGGGGAGAACCGTATATAAGTGCAGTAGTCGCCGTGAACGTTCTTTTTCGCAACGGGTTTGCCGCCAGAACACAGGTAAGTGCCGTGTG
TGGTTCCCGCGGGCCTGGCCTCTTTACGGGTTATGGCCCTTGCGTGCCTTGAATTACTTCCACTGGCTGCAGTACGTGATTCTTGATCCCGAGCTTCGGGTTGGAAGTGGGTGG
GAGAGTTCGAGGCCTTGCGCTTAAGGAGCCCCTTCGCCTCGTGCTTGAGTTGAGGCCTGGCCTGGGCGCTGGGGCCGCCGCGTGCGAATCTGGTGGCACCTTCGCGCCTGTCTC
GCTGCTTTCGATAAGTCTCTAGCCATTTAAAATTTTTGATGACCTGCTGCGACGCTTTTTTTCTGGCAAGATAGTCTTGTAAATGCGGGCCAAGATCTGCACACTGGTATTTCG
GTTTTTGGGGCCGCGGGCGGCGACGGGGCCCGTGCGTCCCAGCGCACATGTTCGGCGAGGCGGGGCCTGCGAGCGCGGCCACCGAGAATCGGACGGGGGTAGTCTCAAGCTGGC
CGGCCTGCTCTGGTGCCTGGCCTCGCGCCGCCGTGTATCGCCCCGCCCTGGGCGGCAAGGCTGGCCCGGTCGGCACCAGTTGCGTGAGCGGAAAGATGGCCGCTTCCCGGCCCT
GCTGCAGGGAGCTCAAAATGGAGGACGCGGCGCTCGGGAGAGCGGGCGGGTGAGTCACCCACACAAAGGAAAAGGGCCTTTCCGTCCTCAGCCGTCGCTTCATGTGACTCCACG
GAGTACCGGGCGCCGTCCAGGCACCTCGATTAGTTCTCGAGCTTTTGGAGTACGTCGTCTTTAGGTTGGGGGGAGGGGTTTTATGCGATGGAGTTTCCCCACACTGAGTGGGTG
GAGACTGAAGTTAGGCCAGCTTGGCACTTGATGTAATTCTCCTTGGAATTTGCCCTTTTTGAGTTTGGATCTTGGTTCATTCTCAAGCCTCAGACAGTGGTTCAAAGTTTTTTT
CTTCCATTTCAGGTGTCGTGAGGAATTCTGCAGTCGATCGACGGTACCGCGGGCCCGGGGCTCGAGATCTAGAGTCGAGAAGCTTGATGATCTGCGCAGCACCATGGCCTGAAA
TAACCTCTGAAAGAGGAACTTGGTTAGGTACCTTCTGAGGCGGAAAGAACCAGCTGTGGAATGTGTGTCAGTTAGGGTGTGGAAAGTCCCCAGGCTCCCCAGCAGGCAGAAGTA
TGCAAAGCATGCATCTCAATTAGTCAGCAACCAGGTGTGGAAAGTCCCCAGGCTCCCCAGCAGGCAGAAGTATGCAAAGCATGCATCTCAATTAGTCAGCAACCATAGTCCCGC
CCCTAACTCCGCCCATCCCGCCCCTAACTCCGCCCAGTTCCGCCCATTCTCCGCCCCATGGCTGACTAATTTTTTTTATTTATGCAGAGGCCGAGGCCGCCTCGGCCTCTGAGC
TATTCCAGAAGTAGTGAGGAGGCTTTTTTGGAGGCCTAGGCTTTTGCAAAAAGCTTACCATGACCGAGTACAAGCCCACGGTGCGCCTCGCCACCCGCGACGACGTCCCCAGGG
CCGTACGCACCCTCGCCGCCGCGTTCGCCGACTACCCCGCCACGCGCCACACCGTCGATCCGGACCGCCACATCGAGCGGGTCACCGAGCTGCAAGAACTCTTCCTCACGCGCG
TCGGGCTCGACATCGGCAAGGTGTGGGTCGCGGACGACGGCGCCGCGGTGGCGGTCTGGACCACGCCGGAGAGCGTCGAAGCGGGGGCGGTGTTCGCCGAGATCGGCCCGCGCA
TGGCCGAGTTGAGCGGTTCCCGGCTGGCCGCGCAGCAACAGATGGAAGGCCTCCTGGCGCCGCACCGGCCCAAGGAGCCCGCGTGGTTCCTGGCCACCGTCGGCGTCTCGCCCG
ACCACCAGGGCAAGGGTCTGGGCAGCGCCGTCGTGCTCCCCGGAGTGGAGGCGGCCGAGCGCGCCGGGGTGCCCGCCTTCCTGGAGACCTCCGCGCCCCGCAACCTCCCCTTCT
ACGAGCGGCTCGGCTTCACCGTCACCGCCGACGTCGAGTGCCCGAAGGACCGCGCGACCTGGTGCATGACCCGCAAGCCCGGTGCCTGACGCCCGCCCCACGACCCGCAGCGCC
CGACCGAAAGGAGCGCACGACCCCATGCCAGTCGACAATCAACCTCTGGATTACAAAATTTGTGAAAGATTGACTGGTATTCTTAACTATGTTGCTCCTTTTACGCTATGTGGA
TACGCTGCTTTAATGCCTTTGTATCATGCTATTGCTTCCCGTATGGCTTTCATTTTCTCCTCCTTGTATAAATCCTGGTTGCTGTCTCTTTATGAGGAGTTGTGGCCCGTTGTC
AGGCAACGTGGCGTGGTGTGCACTGTGTTTGCTGACGCAACCCCCACTGGTTGGGGCATTGCCACCACCTGTCAGCTCCTTTCCGGGACTTTCGCTTTCCCCCTCCCTATTGCC
ACGGCGGAACTCATCGCCGCCTGCCTTGCCCGCTGCTGGACAGGGGCTCGGCTGTTGGGCACTGACAATTCCGTGGTGTTGTCGGGGAAGCTGACGTCCTTTCCATGGCTGCTC
GCCTGTGTTGCCACCTGGATTCTGCGCGGGACGTCCTTCTGCTACGTCCCTTCGGCCCTCAATCCAGCGGACCTTCCTTCCCGCGGCCTGCTGCCGGCTCTGCGGCCTCTTCCG
CGTCTTCGCCTTCGCCCTCAGACGAGTCGGATCTCCCTTTGGGCCGCCTCCCCGCCTGGAATTCGAGCTCGGTACCTTTAAGACCAATGACTTACAAGGCAGCTGTAGATCTTA
GCCACTTTTTAAAAGAAAAGGGGGGACTGGAAGGGCTAATTCACTCCCAACGAAGACAAGATCTGCTTTTTGCTTGTACTGGGTCTCTCTGGTTAGACCAGATCTGAGCCTGGG
AGCTCTCTGGCTAACTAGGGAACCCACTGCTTAAGCCTCAATAAAGCTTGCCTTGAGTGCTTCAAGTAGTGTGTGCCCGTCTGTTGTGTGACTCTGGTAACTAGAGATCCCTCA
GACCCTTTTAGTCAGTGTGGAAAATCTCTAGCAGTAGTAGTTCATGTCATCTTATTATTCAGTATTTATAACTTGCAAAGAAATGAATATCAGAGAGTGAGAGGAACTTGTTTA
TTGCAGCTTATAATGGTTACAAATAAAGCAATAGCATCACAAATTTCACAAATAAAGCATTTTTTTCACTGCATTCTAGTTGTGGTTTGTCCAAACTCATCAATGTATCTTATC
ATGTCTGGCTCTAGCTATCCCGCCCCTAACTCCGCCCAGTTCCGCCCATTCTCCGCCCCATGGCTGACTAATTTTTTTTATTTATGCAGAGGCCGAGGCCGCCTCGGCCTCTGA
GCTATTCCAGAAGTAGTGAGGAGGCTTTTTTGGAGGCCTAGGCTTTTGCGTCGAGACGTACCCAATTCGCCCTATAGTGAGTCGTATTACGCGCGCTCACTGGCCGTCGTTTTA
CAACGTCGTGACTGGGAAAACCCTGGCGTTACCCAACTTAATCGCCTTGCAGCACATCCCCCTTTCGCCAGCTGGCGTAATAGCGAAGAGGCCCGCACCGATCGCCCTTCCCAA
CAGTTGCGCAGCCTGAATGGCGAATGGCGCGACGCGCCCTGTAGCGGCGCATTAAGCGCGGCGGGTGTGGTGGTTACGCGCAGCGTGACCGCTACACTTGCCAGCGCCCTAGCG
CCCGCTCCTTTCGCTTTCTTCCCTTCCTTTCTCGCCACGTTCGCCGGCTTTCCCCGTCAAGCTCTAAATCGGGGGCTCCCTTTAGGGTTCCGATTTAGTGCTTTACGGCACCTC
GACCCCAAAAAACTTGATTAGGGTGATGGTTCACGTAGTGGGCCATCGCCCTGATAGACGGTTTTTCGCCCTTTGACGTTGGAGTCCACGTTCTTTAATAGTGGACTCTTGTTC
CAAACTGGAACAACACTCAACCCTATCTCGGTCTATTCTTTTGATTTATAAGGGATTTTGCCGATTTCGGCCTATTGGTTAAAAAATGAGCTGATTTAACAAAAATTTAACGCG
AATTTTAACAAAATATTAACGTTTACAATTTCCCAGGTGGCACTTTTCGGGGAAATGTGCGCGGAACCCCTATTTGTTTATTTTTCTAAATACATTCAAATATGTATCCGCTCA
TGAGACAATAACCCTGATAAATGCTTCAATAATATTGAAAAAGGAAGAGTATGAGTATTCAACATTTCCGTGTCGCCCTTATTCCCTTTTTTGCGGCATTTTGCCTTCCTGTTT
TTGCTCACCCAGAAACGCTGGTGAAAGTAAAAGATGCTGAAGATCAGTTGGGTGCACGAGTGGGTTACATCGAACTGGATCTCAACAGCGGTAAGATCCTTGAGAGTTTTCGCC
CCGAAGAACGTTTTCCAATGATGAGCACTTTTAAAGTTCTGCTATGTGGCGCGGTATTATCCCGTATTGACGCCGGGCAAGAGCAACTCGGTCGCCGCATACACTATTCTCAGA
ATGACTTGGTTGAGTACTCACCAGTCACAGAAAAGCATCTTACGGATGGCATGACAGTAAGAGAATTATGCAGTGCTGCCATAACCATGAGTGATAACACTGCGGCCAACTTAC
TTCTGACAACGATCGGAGGACCGAAGGAGCTAACCGCTTTTTTGCACAACATGGGGGATCATGTAACTCGCCTTGATCGTTGGGAACCGGAGCTGAATGAAGCCATACCAAACG

ACGAGCGTGACACCACGATGCCTGTAGCAATGGCAACAACGTTGCGCAAACTATTAACTGGCGAACTACTTACTCTAGCTTCCCGGCAACAATTAATAGACTGGATGGAGGCGG
ATAAAGTTGCAGGACCACTTCTGCGCTCGGCCCTTCCGGCTGGCTGGTTTATTGCTGATAAATCTGGAGCCGGTGAGCGTGGGTCTCGCGGTATCATTGCAGCACTGGGGCCAG
ATGGTAAGCCCTCCCGTATCGTAGTTATCTACACGACGGGGAGTCAGGCAACTATGGATGAACGAAATAGACAGATCGCTGAGATAGGTGCCTCACTGATTAAGCATTGGTAAC
TGTCAGACCAAGTTTACTCATATATACTTTAGATTGATTTAAAACTTCATTTTTAATTTAAAAGGATCTAGGTGAAGATCCTTTTTGATAATCTCATGACCAAAATCCCTTAAC
GTGAGTTTTCGTTCCACTGAGCGTCAGACCCCGTAGAAAAGATCAAAGGATCTTCTTGAGATCCTTTTTTTCTGCGCGTAATCTGCTGCTTGCAAACAAAAAAACCACCGCTAC
CAGCGGTGGTTTGTTTGCCGGATCAAGAGCTACCAACTCTTTTTCCGAAGGTAACTGGCTTCAGCAGAGCGCAGATACCAAATACTGTCCTTCTAGTGTAGCCGTAGTTAGGCC
ACCACTTCAAGAACTCTGTAGCACCGCCTACATACCTCGCTCTGCTAATCCTGTTACCAGTGGCTGCTGCCAGTGGCGATAAGTCGTGTCTTACCGGGTTGGACTCAAGACGAT
AGTTACCGGATAAGGCGCAGCGGTCGGGCTGAACGGGGGGTTCGTGCACACAGCCCAGCTTGGAGCGAACGACCTACACCGAACTGAGATACCTACAGCGTGAGCTATGAGAAA
GCGCCACGCTTCCCGAAGGGAGAAAGGCGGACAGGTATCCGGTAAGCGGCAGGGTCGGAACAGGAGAGCGCACGAGGGAGCTTCCAGGGGGAAACGCCTGGTATCTTTATAGTC
CTGTCGGGTTTCGCCACCTCTGACTTGAGCGTCGATTTTTGTGATGCTCGTCAGGGGGGCGGAGCCTATGGAAAAACGCCAGCAACGCGGCCTTTTTACGGTTCCTGGCCTTTT
GCTGGCCTTTTGCTCACATGTTCTTTCCTGCGTTATCCCCTGATTCTGTGGATAACCGTATTACCGCCTTTGAGTGAGCTGATACCGCTCGCCGCAGCCGAACGACCGAGCGCA
GCGAGTCAGTGAGCGAGGAAGCGGAAGAGCGCCCAATACGCAAACCGCCTCTCCCCGCGCGTTGGCCGATTCATTAATGCAGCTGGCACGACAGGTTTCCCGACTGGAAAGCGG
GCAGTGAGCGCAACGCAATTAATGTGAGTTAGCTCACTCATTAGGCACCCCAGGCTTTACACTTTATGCTTCCGGCTCGTATGTTGTGTGGAATTGTGAGCGGATAACAATTTC
ACACAGGAAACAGCTATGACCATGATTACGCCAAGCGCGCAATTAACCCTCACTAAAGGGAACAAAAGCTGGAGCTGCAAGCTTAATGTAGTCTTATGCAATACTCTTGTAGTC
TTGCAACATGGTAACGATGAGTTAGCAACATGCCTTACAAGGAGAGAAAAAGCACCGTGCATGCCGATTGGTGGAAGTAAGGTGGTACGATCGTGCCTTATTAGGAAGGCAACA
GACGGGTCTGACATGGATTGGACGAACCACTGAATTGCCGCATTGCAGAGATATTGTATTTAAGTGCCTAGCTCGATACAATAAACGGGTCTCTCTGGTTAGACCAGATCTGAG
CCTGGGAGCTCTCTGGCTAACTAGGGAACCCACTGCTTAAGCCTCAATAAAGCTTGCCTTGAGTGCTTCAAGTAGTGTGTGCCCGTCTGTTGTGTGACTCTGGTAACTAGAGAT
CCCTCAGACCCTTTTAGTCAGTGTGGAAAATCTCTAGCAGTGGCGCCCGAACAGGGACCTGAAAGCGAAAGGGAAACCAGAGCTCTCTCGACGCAGGACTCGGCTTGCTGAAGC
GCGCACGGCAAGAGGCGAGGGGCGGCGACTGGTGAGTACGCCAAAAATTTTGACTAGCGGAGGCTAGAAGGAGAGAGATGGGTGCGAGAGCGTCAGTATTAAGCGGGGGAGAAT
TAGATCGCGATGGGAAAAAATTCGGTTAAGGCCAGGGGGAAAGAAAAAATATAAATTAAAACATATAGTATGGGCAAGCAGGGAGCTAGAACGATTCGCAGTTAATCCTGGCCT
GTTAGAAACATCAGAAGGCTGTAGACAAATACTGGGACAGCTACAACCATCCCTTCAGACAGGATCAGAAGAACTTAGATCATTATATAATACAGTAGCAACCCTCTATTGTGT
GCATCAAAGGATAGAGATAAAAGACACCAAGGAAGCTTTAGACAAGATAGAGGAAGAGCAAAACAAAAGTAAGACCACCGCACAGCAAGCGGCCGCTGATCTTCAGACCTGGAG
GAGGAGATATGAGGGACAATTGGAGAAGTGAATTATATAAATATAAAGTAGTAAAAATTGAACCATTAGGAGTAGCACCCACCAAGGCAAAGAGAAGAGTGGTGCAGAGAGAAA
AAAGAGCAGTGGGAATAGGAGCTTTGTTCCTTGGGTTCTTGGGAGCAGCAGGAAGCACTATGGGCGCAGCCTCAATGACGCTGACGGTACAGGCCAGACAATTATTGTCTGGTA
TAGTGCAGCAGCAGAACAATTTGCTGAGGGCTATTGAGGCGCAACAGCATCTGTTGCAACTCACAGTCTGGGGCATCAAGCAGCTCCAGGCAAGAATCCTGGCTGTGGAAAGAT
ACCTAAAGGATCAACAGCTCCTGGGGATTTGGGGTTGCTCTGGAAAACTCATTTGCACCACTGCTGTGCCTTGGAATGCTAGTTGGAGTAATAAATCTCTGGAACAGATTTGGA
ATCACACGACCTGGATGGAGTGGGACAGAGAAATTAACAATTACACAAGCTTAATACACTCCTTAATTGAAGAATCGCAAAACCAGCAAGAAAAGAATGAACAAGAATTATTGG
AATTAGATAAATGGGCAAGTTTGTGGAATTGGTTTAACATAACAAATTGGCTGTGGTATATAAAATTATTCATAATGATAGTAGGAGGCTTGGTAGGTTTAAGAATAGTTTTTG
CTGTACTTTCTATAGTGAATAGAGTTAGGCAGGGATATTCACCATTATCGTTTCAGACCCACCTCCCAACCCCGAGGGGACCCGACAGGCCCGAAGGAATAGAAGAAGAAGGTG
GAGAGAGAGACAGAGACAGATCCATTCGATTAGTGAACGGATCTCGACGGTATCGGTTAACTTTTAAAAGAAAAGGGGGGATTGGGGGGTACAGTGCAGGGGAAAGAATAGTAG
ACATAATAGCAACAGACATACAAACTAAAGAATTACAAAAACAAATTACAAAAATTCAAAATTTTATCGATCACGAGACTAGCCTC

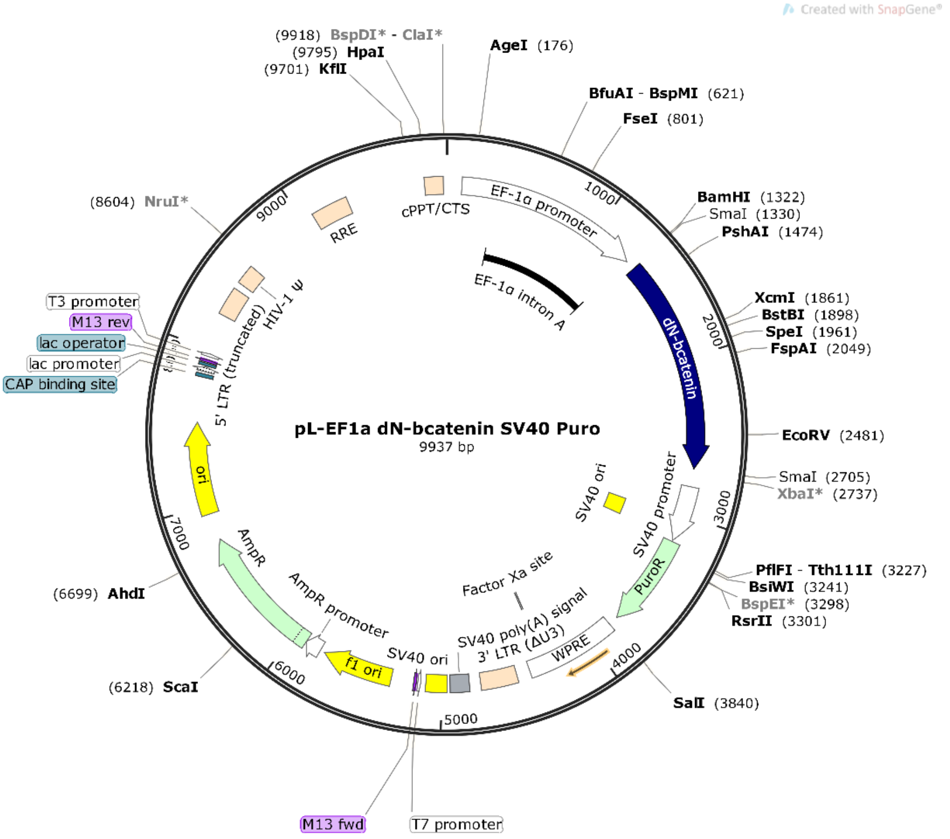

### >pL-EF1a ΔN-βcatenin SV40 Puro

CTAGCCCCGATAAGCTTTGCAAAGATGGATAAAGTTTTAAACAGAGAGGAATCTTTGCAGCTAATGGACCTTCTAGGTCTTGAAAGGAGTGGGAATTGGCTCCGGTGCCCGTCA
GTGGGCAGAGCGCACATCGCCCACAGTCCCCGAGAAGTTGGGGGGAGGGGTCGGCAATTGAACCGGTGCCTAGAGAAGGTGGCGCGGGGTAAACTGGGAAAGTGATGTCGTGTA
CTGGCTCCGCCTTTTTCCCGAGGGTGGGGGAGAACCGTATATAAGTGCAGTAGTCGCCGTGAACGTTCTTTTTCGCAACGGGTTTGCCGCCAGAACACAGGTAAGTGCCGTGTG
TGGTTCCCGCGGGCCTGGCCTCTTTACGGGTTATGGCCCTTGCGTGCCTTGAATTACTTCCACTGGCTGCAGTACGTGATTCTTGATCCCGAGCTTCGGGTTGGAAGTGGGTGG
GAGAGTTCGAGGCCTTGCGCTTAAGGAGCCCCTTCGCCTCGTGCTTGAGTTGAGGCCTGGCCTGGGCGCTGGGGCCGCCGCGTGCGAATCTGGTGGCACCTTCGCGCCTGTCTC
GCTGCTTTCGATAAGTCTCTAGCCATTTAAAATTTTTGATGACCTGCTGCGACGCTTTTTTTCTGGCAAGATAGTCTTGTAAATGCGGGCCAAGATCTGCACACTGGTATTTCG
GTTTTTGGGGCCGCGGGCGGCGACGGGGCCCGTGCGTCCCAGCGCACATGTTCGGCGAGGCGGGGCCTGCGAGCGCGGCCACCGAGAATCGGACGGGGGTAGTCTCAAGCTGGC
CGGCCTGCTCTGGTGCCTGGCCTCGCGCCGCCGTGTATCGCCCCGCCCTGGGCGGCAAGGCTGGCCCGGTCGGCACCAGTTGCGTGAGCGGAAAGATGGCCGCTTCCCGGCCCT
GCTGCAGGGAGCTCAAAATGGAGGACGCGGCGCTCGGGAGAGCGGGCGGGTGAGTCACCCACACAAAGGAAAAGGGCCTTTCCGTCCTCAGCCGTCGCTTCATGTGACTCCACG
GAGTACCGGGCGCCGTCCAGGCACCTCGATTAGTTCTCGAGCTTTTGGAGTACGTCGTCTTTAGGTTGGGGGGAGGGGTTTTATGCGATGGAGTTTCCCCACACTGAGTGGGTG
GAGACTGAAGTTAGGCCAGCTTGGCACTTGATGTAATTCTCCTTGGAATTTGCCCTTTTTGAGTTTGGATCTTGGTTCATTCTCAAGCCTCAGACAGTGGTTCAAAGTTTTTTT
CTTCCATTTCAGGTGTCGTGAGGAATTCTGCAGTCGATCGACGGTACCGCGGGCCCTCCGCGGCCGCGGATCCCCCGGGCCACATGAGGACCTACACTTATGAGAAGCTTCTGT
GGACCACAAGCAGAGTGCTGAAGGTGCTGTCTGTCTGCTCTAGCAACAAGCCGGCCATTGTAGAAGCTGGTGGGATGCAGGCACTGGGGCTTCATCTGACAGACCCAAGTCAGC
GACTTGTTCAAAACTGTCTTTGGACTCTCAGAAACCTTTCAGATGCAGCGACTAAGCAGGAAGGGATGGAAGGCCTCCTTGGGACTCTAGTGCAGCTTCTGGGTTCCGATGATA
TAAATGTGGTCACCTGTGCAGCTGGAATTCTCTCTAACCTCACTTGCAATAATTACAAAAACAAGATGATGGTGTGCCAAGTGGGTGGCATAGAGGCTCTTGTACGCACCGTCC
TTCGTGCTGGTGACAGGGAAGACATCACTGAGCCTGCCATCTGTGCTCTTCGTCATCTGACCAGCCGGCATCAGGAAGCCGAGATGGCCCAGAATGCCGTTCGCCTTCATTATG
GACTGCCTGTTGTGGTTAAACTCCTGCACCCACCATCCCACTGGCCTCTGATAAAGGCAACTGTTGGATTGATTCGAAACCTTGCCCTTTGCCCAGCAAATCATGCGCCTTTGC
GGGAACAGGGTGCTATTCCACGACTAGTTCAGCTGCTTGTACGAGCACATCAGGACACCCAACGGCGCACCTCCATGGGTGGAACGCAGCAGCAGTTTGTGGAGGGCGTGCGCA

TGGAGGAGATAGTAGAAGGGTGTACTGGAGCTCTCCACATCCTTGCTCGGGACGTTCACAACCGGATTGTAATCCGAGGACTCAATACCATTCCATTGTTTGTGCAGTTGCTTT
ATTCTCCCATTGAAAATATCCAAAGAGTAGCTGCAGGGGTCCTCTGTGAACTTGCTCAGGACAAGGAGGCTGCAGAGGCCATTGAAGCTGAGGGAGCCACAGCTCCCCTGACAG
AGTTACTCCACTCCAGGAATGAAGGCGTGGCAACATACGCAGCTGCTGTCCTATTCCGAATGTCTGAGGACAAGCCACAGGATTACAAGAAGCGGCTTTCAGTCGAGCTGACCA
GTTCCCTCTTCAGGACAGAGCCAATGGCTTGGAATGAGACTGCAGATCTTGGACTGGACATTGGTGCCCAGGGAGAAGCCCTTGGATATCGCCAGGATGATCCCAGCTACCGTT
CTTTTCACTCTGGTGGATACGGCCAGGATGCCTTGGGGATGGACCCTATGATGGAGCATGAGATGGGTGGCCACCACCCTGGTGCTGACTATCCAGTTGATGGGCTGCCTGATC
TGGGACACGCCCAGGACCTCATGGATGGGCTGCCCCCAGGTGATAGCAATCAGCTGGCCTGGTTTGATACTGACCTGTAACCCGGGGAATTCCTCGAGAAGCTGGGGCTCGAGA
TCTAGAGTCGAGAAGCTTGATGATCTGCGCAGCACCATGGCCTGAAATAACCTCTGAAAGAGGAACTTGGTTAGGTACCTTCTGAGGCGGAAAGAACCAGCTGTGGAATGTGTG
TCAGTTAGGGTGTGGAAAGTCCCCAGGCTCCCCAGCAGGCAGAAGTATGCAAAGCATGCATCTCAATTAGTCAGCAACCAGGTGTGGAAAGTCCCCAGGCTCCCCAGCAGGCAG
AAGTATGCAAAGCATGCATCTCAATTAGTCAGCAACCATAGTCCCGCCCCTAACTCCGCCCATCCCGCCCCTAACTCCGCCCAGTTCCGCCCATTCTCCGCCCCATGGCTGACT
AATTTTTTTTATTTATGCAGAGGCCGAGGCCGCCTCGGCCTCTGAGCTATTCCAGAAGTAGTGAGGAGGCTTTTTTGGAGGCCTAGGCTTTTGCAAAAAGCTTACCATGACCGA
GTACAAGCCCACGGTGCGCCTCGCCACCCGCGACGACGTCCCCAGGGCCGTACGCACCCTCGCCGCCGCGTTCGCCGACTACCCCGCCACGCGCCACACCGTCGATCCGGACCG
CCACATCGAGCGGGTCACCGAGCTGCAAGAACTCTTCCTCACGCGCGTCGGGCTCGACATCGGCAAGGTGTGGGTCGCGGACGACGGCGCCGCGGTGGCGGTCTGGACCACGCC
GGAGAGCGTCGAAGCGGGGGCGGTGTTCGCCGAGATCGGCCCGCGCATGGCCGAGTTGAGCGGTTCCCGGCTGGCCGCGCAGCAACAGATGGAAGGCCTCCTGGCGCCGCACCG
GCCCAAGGAGCCCGCGTGGTTCCTGGCCACCGTCGGCGTCTCGCCCGACCACCAGGGCAAGGGTCTGGGCAGCGCCGTCGTGCTCCCCGGAGTGGAGGCGGCCGAGCGCGCCGG
GGTGCCCGCCTTCCTGGAGACCTCCGCGCCCCGCAACCTCCCCTTCTACGAGCGGCTCGGCTTCACCGTCACCGCCGACGTCGAGTGCCCGAAGGACCGCGCGACCTGGTGCAT
GACCCGCAAGCCCGGTGCCTGACGCCCGCCCCACGACCCGCAGCGCCCGACCGAAAGGAGCGCACGACCCCATGCCAGTCGACAATCAACCTCTGGATTACAAAATTTGTGAAA
GATTGACTGGTATTCTTAACTATGTTGCTCCTTTTACGCTATGTGGATACGCTGCTTTAATGCCTTTGTATCATGCTATTGCTTCCCGTATGGCTTTCATTTTCTCCTCCTTGT
ATAAATCCTGGTTGCTGTCTCTTTATGAGGAGTTGTGGCCCGTTGTCAGGCAACGTGGCGTGGTGTGCACTGTGTTTGCTGACGCAACCCCCACTGGTTGGGGCATTGCCACCA
CCTGTCAGCTCCTTTCCGGGACTTTCGCTTTCCCCCTCCCTATTGCCACGGCGGAACTCATCGCCGCCTGCCTTGCCCGCTGCTGGACAGGGGCTCGGCTGTTGGGCACTGACA
ATTCCGTGGTGTTGTCGGGGAAGCTGACGTCCTTTCCATGGCTGCTCGCCTGTGTTGCCACCTGGATTCTGCGCGGGACGTCCTTCTGCTACGTCCCTTCGGCCCTCAATCCAG
CGGACCTTCCTTCCCGCGGCCTGCTGCCGGCTCTGCGGCCTCTTCCGCGTCTTCGCCTTCGCCCTCAGACGAGTCGGATCTCCCTTTGGGCCGCCTCCCCGCCTGGAATTCGAG
CTCGGTACCTTTAAGACCAATGACTTACAAGGCAGCTGTAGATCTTAGCCACTTTTTAAAAGAAAAGGGGGGACTGGAAGGGCTAATTCACTCCCAACGAAGACAAGATCTGCT
TTTTGCTTGTACTGGGTCTCTCTGGTTAGACCAGATCTGAGCCTGGGAGCTCTCTGGCTAACTAGGGAACCCACTGCTTAAGCCTCAATAAAGCTTGCCTTGAGTGCTTCAAGT
AGTGTGTGCCCGTCTGTTGTGTGACTCTGGTAACTAGAGATCCCTCAGACCCTTTTAGTCAGTGTGGAAAATCTCTAGCAGTAGTAGTTCATGTCATCTTATTATTCAGTATTT
ATAACTTGCAAAGAAATGAATATCAGAGAGTGAGAGGAACTTGTTTATTGCAGCTTATAATGGTTACAAATAAAGCAATAGCATCACAAATTTCACAAATAAAGCATTTTTTTC
ACTGCATTCTAGTTGTGGTTTGTCCAAACTCATCAATGTATCTTATCATGTCTGGCTCTAGCTATCCCGCCCCTAACTCCGCCCAGTTCCGCCCATTCTCCGCCCCATGGCTGA
CTAATTTTTTTTATTTATGCAGAGGCCGAGGCCGCCTCGGCCTCTGAGCTATTCCAGAAGTAGTGAGGAGGCTTTTTTGGAGGCCTAGGCTTTTGCGTCGAGACGTACCCAATT
CGCCCTATAGTGAGTCGTATTACGCGCGCTCACTGGCCGTCGTTTTACAACGTCGTGACTGGGAAAACCCTGGCGTTACCCAACTTAATCGCCTTGCAGCACATCCCCCTTTCG
CCAGCTGGCGTAATAGCGAAGAGGCCCGCACCGATCGCCCTTCCCAACAGTTGCGCAGCCTGAATGGCGAATGGCGCGACGCGCCCTGTAGCGGCGCATTAAGCGCGGCGGGTG
TGGTGGTTACGCGCAGCGTGACCGCTACACTTGCCAGCGCCCTAGCGCCCGCTCCTTTCGCTTTCTTCCCTTCCTTTCTCGCCACGTTCGCCGGCTTTCCCCGTCAAGCTCTAA
ATCGGGGGCTCCCTTTAGGGTTCCGATTTAGTGCTTTACGGCACCTCGACCCCAAAAAACTTGATTAGGGTGATGGTTCACGTAGTGGGCCATCGCCCTGATAGACGGTTTTTC
GCCCTTTGACGTTGGAGTCCACGTTCTTTAATAGTGGACTCTTGTTCCAAACTGGAACAACACTCAACCCTATCTCGGTCTATTCTTTTGATTTATAAGGGATTTTGCCGATTT
CGGCCTATTGGTTAAAAAATGAGCTGATTTAACAAAAATTTAACGCGAATTTTAACAAAATATTAACGTTTACAATTTCCCAGGTGGCACTTTTCGGGGAAATGTGCGCGGAAC
CCCTATTTGTTTATTTTTCTAAATACATTCAAATATGTATCCGCTCATGAGACAATAACCCTGATAAATGCTTCAATAATATTGAAAAAGGAAGAGTATGAGTATTCAACATTT
CCGTGTCGCCCTTATTCCCTTTTTTGCGGCATTTTGCCTTCCTGTTTTTGCTCACCCAGAAACGCTGGTGAAAGTAAAAGATGCTGAAGATCAGTTGGGTGCACGAGTGGGTTA
CATCGAACTGGATCTCAACAGCGGTAAGATCCTTGAGAGTTTTCGCCCCGAAGAACGTTTTCCAATGATGAGCACTTTTAAAGTTCTGCTATGTGGCGCGGTATTATCCCGTAT
TGACGCCGGGCAAGAGCAACTCGGTCGCCGCATACACTATTCTCAGAATGACTTGGTTGAGTACTCACCAGTCACAGAAAAGCATCTTACGGATGGCATGACAGTAAGAGAATT
ATGCAGTGCTGCCATAACCATGAGTGATAACACTGCGGCCAACTTACTTCTGACAACGATCGGAGGACCGAAGGAGCTAACCGCTTTTTTGCACAACATGGGGGATCATGTAAC
TCGCCTTGATCGTTGGGAACCGGAGCTGAATGAAGCCATACCAAACGACGAGCGTGACACCACGATGCCTGTAGCAATGGCAACAACGTTGCGCAAACTATTAACTGGCGAACT
ACTTACTCTAGCTTCCCGGCAACAATTAATAGACTGGATGGAGGCGGATAAAGTTGCAGGACCACTTCTGCGCTCGGCCCTTCCGGCTGGCTGGTTTATTGCTGATAAATCTGG
AGCCGGTGAGCGTGGGTCTCGCGGTATCATTGCAGCACTGGGGCCAGATGGTAAGCCCTCCCGTATCGTAGTTATCTACACGACGGGGAGTCAGGCAACTATGGATGAACGAAA
TAGACAGATCGCTGAGATAGGTGCCTCACTGATTAAGCATTGGTAACTGTCAGACCAAGTTTACTCATATATACTTTAGATTGATTTAAAACTTCATTTTTAATTTAAAAGGAT
CTAGGTGAAGATCCTTTTTGATAATCTCATGACCAAAATCCCTTAACGTGAGTTTTCGTTCCACTGAGCGTCAGACCCCGTAGAAAAGATCAAAGGATCTTCTTGAGATCCTTT
TTTTCTGCGCGTAATCTGCTGCTTGCAAACAAAAAAACCACCGCTACCAGCGGTGGTTTGTTTGCCGGATCAAGAGCTACCAACTCTTTTTCCGAAGGTAACTGGCTTCAGCAG
AGCGCAGATACCAAATACTGTCCTTCTAGTGTAGCCGTAGTTAGGCCACCACTTCAAGAACTCTGTAGCACCGCCTACATACCTCGCTCTGCTAATCCTGTTACCAGTGGCTGC
TGCCAGTGGCGATAAGTCGTGTCTTACCGGGTTGGACTCAAGACGATAGTTACCGGATAAGGCGCAGCGGTCGGGCTGAACGGGGGGTTCGTGCACACAGCCCAGCTTGGAGCG
AACGACCTACACCGAACTGAGATACCTACAGCGTGAGCTATGAGAAAGCGCCACGCTTCCCGAAGGGAGAAAGGCGGACAGGTATCCGGTAAGCGGCAGGGTCGGAACAGGAGA
GCGCACGAGGGAGCTTCCAGGGGGAAACGCCTGGTATCTTTATAGTCCTGTCGGGTTTCGCCACCTCTGACTTGAGCGTCGATTTTTGTGATGCTCGTCAGGGGGGCGGAGCCT
ATGGAAAAACGCCAGCAACGCGGCCTTTTTACGGTTCCTGGCCTTTTGCTGGCCTTTTGCTCACATGTTCTTTCCTGCGTTATCCCCTGATTCTGTGGATAACCGTATTACCGC
CTTTGAGTGAGCTGATACCGCTCGCCGCAGCCGAACGACCGAGCGCAGCGAGTCAGTGAGCGAGGAAGCGGAAGAGCGCCCAATACGCAAACCGCCTCTCCCCGCGCGTTGGCC
GATTCATTAATGCAGCTGGCACGACAGGTTTCCCGACTGGAAAGCGGGCAGTGAGCGCAACGCAATTAATGTGAGTTAGCTCACTCATTAGGCACCCCAGGCTTTACACTTTAT
GCTTCCGGCTCGTATGTTGTGTGGAATTGTGAGCGGATAACAATTTCACACAGGAAACAGCTATGACCATGATTACGCCAAGCGCGCAATTAACCCTCACTAAAGGGAACAAAA
GCTGGAGCTGCAAGCTTAATGTAGTCTTATGCAATACTCTTGTAGTCTTGCAACATGGTAACGATGAGTTAGCAACATGCCTTACAAGGAGAGAAAAAGCACCGTGCATGCCGA
TTGGTGGAAGTAAGGTGGTACGATCGTGCCTTATTAGGAAGGCAACAGACGGGTCTGACATGGATTGGACGAACCACTGAATTGCCGCATTGCAGAGATATTGTATTTAAGTGC
CTAGCTCGATACAATAAACGGGTCTCTCTGGTTAGACCAGATCTGAGCCTGGGAGCTCTCTGGCTAACTAGGGAACCCACTGCTTAAGCCTCAATAAAGCTTGCCTTGAGTGCT
TCAAGTAGTGTGTGCCCGTCTGTTGTGTGACTCTGGTAACTAGAGATCCCTCAGACCCTTTTAGTCAGTGTGGAAAATCTCTAGCAGTGGCGCCCGAACAGGGACCTGAAAGCG
AAAGGGAAACCAGAGCTCTCTCGACGCAGGACTCGGCTTGCTGAAGCGCGCACGGCAAGAGGCGAGGGGCGGCGACTGGTGAGTACGCCAAAAATTTTGACTAGCGGAGGCTAG
AAGGAGAGAGATGGGTGCGAGAGCGTCAGTATTAAGCGGGGGAGAATTAGATCGCGATGGGAAAAAATTCGGTTAAGGCCAGGGGGAAAGAAAAAATATAAATTAAAACATATA
GTATGGGCAAGCAGGGAGCTAGAACGATTCGCAGTTAATCCTGGCCTGTTAGAAACATCAGAAGGCTGTAGACAAATACTGGGACAGCTACAACCATCCCTTCAGACAGGATCA
GAAGAACTTAGATCATTATATAATACAGTAGCAACCCTCTATTGTGTGCATCAAAGGATAGAGATAAAAGACACCAAGGAAGCTTTAGACAAGATAGAGGAAGAGCAAAACAAA
AGTAAGACCACCGCACAGCAAGCGGCCGCTGATCTTCAGACCTGGAGGAGGAGATATGAGGGACAATTGGAGAAGTGAATTATATAAATATAAAGTAGTAAAAATTGAACCATT
AGGAGTAGCACCCACCAAGGCAAAGAGAAGAGTGGTGCAGAGAGAAAAAAGAGCAGTGGGAATAGGAGCTTTGTTCCTTGGGTTCTTGGGAGCAGCAGGAAGCACTATGGGCGC
AGCCTCAATGACGCTGACGGTACAGGCCAGACAATTATTGTCTGGTATAGTGCAGCAGCAGAACAATTTGCTGAGGGCTATTGAGGCGCAACAGCATCTGTTGCAACTCACAGT
CTGGGGCATCAAGCAGCTCCAGGCAAGAATCCTGGCTGTGGAAAGATACCTAAAGGATCAACAGCTCCTGGGGATTTGGGGTTGCTCTGGAAAACTCATTTGCACCACTGCTGT
GCCTTGGAATGCTAGTTGGAGTAATAAATCTCTGGAACAGATTTGGAATCACACGACCTGGATGGAGTGGGACAGAGAAATTAACAATTACACAAGCTTAATACACTCCTTAAT
TGAAGAATCGCAAAACCAGCAAGAAAAGAATGAACAAGAATTATTGGAATTAGATAAATGGGCAAGTTTGTGGAATTGGTTTAACATAACAAATTGGCTGTGGTATATAAAATT
ATTCATAATGATAGTAGGAGGCTTGGTAGGTTTAAGAATAGTTTTTGCTGTACTTTCTATAGTGAATAGAGTTAGGCAGGGATATTCACCATTATCGTTTCAGACCCACCTCCC
AACCCCGAGGGGACCCGACAGGCCCGAAGGAATAGAAGAAGAAGGTGGAGAGAGAGACAGAGACAGATCCATTCGATTAGTGAACGGATCTCGACGGTATCGGTTAACTTTTAA
AAGAAAAGGGGGGATTGGGGGGTACAGTGCAGGGGAAAGAATAGTAGACATAATAGCAACAGACATACAAACTAAAGAATTACAAAAACAAATTACAAAAATTCAAAATTTTAT
CGATCACGAGACTAGCCTC

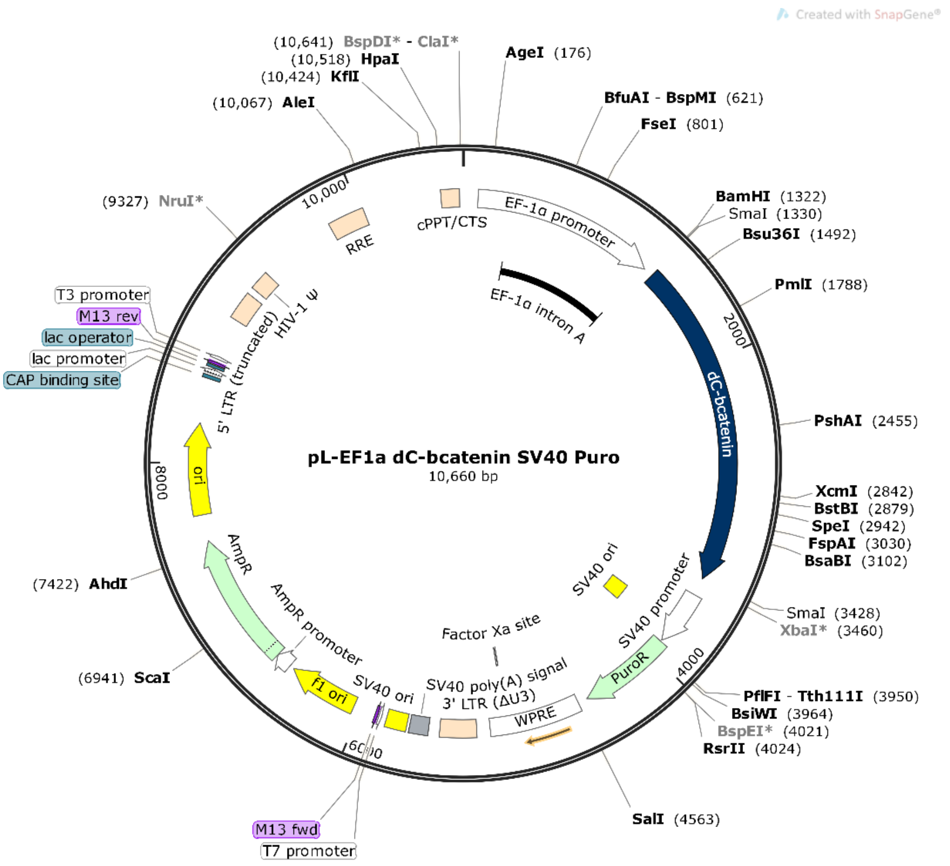

**>pL-EF1a ΔC-βcatenin SV40 Puro**

CTAGCCCCGATAAGCTTTGCAAAGATGGATAAAGTTTTAAACAGAGAGGAATCTTTGCAGCTAATGGACCTTCTAGGTCTTGAAAGGAGTGGGAATTGGCTCCGGTGCCCGTCA
GTGGGCAGAGCGCACATCGCCCACAGTCCCCGAGAAGTTGGGGGGAGGGGTCGGCAATTGAACCGGTGCCTAGAGAAGGTGGCGCGGGGTAAACTGGGAAAGTGATGTCGTGTA
CTGGCTCCGCCTTTTTCCCGAGGGTGGGGGAGAACCGTATATAAGTGCAGTAGTCGCCGTGAACGTTCTTTTTCGCAACGGGTTTGCCGCCAGAACACAGGTAAGTGCCGTGTG
TGGTTCCCGCGGGCCTGGCCTCTTTACGGGTTATGGCCCTTGCGTGCCTTGAATTACTTCCACTGGCTGCAGTACGTGATTCTTGATCCCGAGCTTCGGGTTGGAAGTGGGTGG
GAGAGTTCGAGGCCTTGCGCTTAAGGAGCCCCTTCGCCTCGTGCTTGAGTTGAGGCCTGGCCTGGGCGCTGGGGCCGCCGCGTGCGAATCTGGTGGCACCTTCGCGCCTGTCTC
GCTGCTTTCGATAAGTCTCTAGCCATTTAAAATTTTTGATGACCTGCTGCGACGCTTTTTTTCTGGCAAGATAGTCTTGTAAATGCGGGCCAAGATCTGCACACTGGTATTTCG
GTTTTTGGGGCCGCGGGCGGCGACGGGGCCCGTGCGTCCCAGCGCACATGTTCGGCGAGGCGGGGCCTGCGAGCGCGGCCACCGAGAATCGGACGGGGGTAGTCTCAAGCTGGC
CGGCCTGCTCTGGTGCCTGGCCTCGCGCCGCCGTGTATCGCCCCGCCCTGGGCGGCAAGGCTGGCCCGGTCGGCACCAGTTGCGTGAGCGGAAAGATGGCCGCTTCCCGGCCCT
GCTGCAGGGAGCTCAAAATGGAGGACGCGGCGCTCGGGAGAGCGGGCGGGTGAGTCACCCACACAAAGGAAAAGGGCCTTTCCGTCCTCAGCCGTCGCTTCATGTGACTCCACG
GAGTACCGGGCGCCGTCCAGGCACCTCGATTAGTTCTCGAGCTTTTGGAGTACGTCGTCTTTAGGTTGGGGGGAGGGGTTTTATGCGATGGAGTTTCCCCACACTGAGTGGGTG
GAGACTGAAGTTAGGCCAGCTTGGCACTTGATGTAATTCTCCTTGGAATTTGCCCTTTTTGAGTTTGGATCTTGGTTCATTCTCAAGCCTCAGACAGTGGTTCAAAGTTTTTTT
CTTCCATTTCAGGTGTCGTGAGGAATTCTGCAGTCGATCGACGGTACCGCGGGCCCTCCGCGGCCGCGGATCCCCCGGGCCACATGGCTACTCAAGCTGACCTGATGGAGTTGG
ACATGGCCATGGAGCCGGACAGAAAAGCTGCTGTCAGCCACTGGCAGCAGCAGTCTTACTTGGATTCTGGAATCCATTCTGGTGCCACCACCACAGCTCCTTCCCTGAGTGGCA
AGGGCAACCCTGAGGAAGAAGATGTTGACACCTCCCAAGTCCTTTATGAATGGGAGCAAGGCTTTTCCCAGTCCTTCACGCAAGAGCAAGTAGCTGATATTGACGGGCAGTATG
CAATGACTAGGGCTCAGAGGGTCCGAGCTGCCATGTTCCCTGAGACGCTAGATGAGGGCATGCAGATCCCATCCACGCAGTTTGACGCTGCTCATCCCACTAATGTCCAGCGCT
TGGCTGAACCATCACAGATGTTGAAACATGCAGTTGTCAATTTGATTAACTATCAGGATGACGCGGAACTTGCCACACGTGCAATTCCTGAGCTGACAAAACTGCTAAACGATG
AGGACCAGGTGGTAGTTAATAAAGCTGCTGTTATGGTCCATCAGCTTTCCAAAAAGGAAGCTTCCAGACATGCCATCATGCGCTCCCCTCAGATGGTGTCTGCCATTGTACGCA
CCATGCAGAATACAAATGATGTAGAGACAGCTCGTTGTACTGCTGGGACTCTGCACAACCTTTCTCACCACCGCGAGGGCTTGCTGGCCATCTTTAAGTCTGGTGGCATCCCAG
CGCTGGTGAAAATGCTTGGGTCACCAGTGGATTCTGTACTGTTCTACGCCATCACGACACTGCATAATCTCCTGCTCCATCAGGAAGGAGCTAAAATGGCAGTGCGCCTAGCTG
GTGGACTGCAGAAAATGGTTGCTTTGCTCAACAAAACAAACGTGAAATTCTTGGCTATTACAACAGACTGCCTTCAGATCTTAGCTTATGGCAATCAAGAGAGCAAGCTCATCA
TTCTGGCCAGTGGTGGACCCCAAGCCTTAGTAAACATAATGAGGACCTACACTTATGAGAAGCTTCTGTGGACCACAAGCAGAGTGCTGAAGGTGCTGTCTGTCTGCTCTAGCA
ACAAGCCGGCCATTGTAGAAGCTGGTGGGATGCAGGCACTGGGGCTTCATCTGACAGACCCAAGTCAGCGACTTGTTCAAAACTGTCTTTGGACTCTCAGAAACCTTTCAGATG
CAGCGACTAAGCAGGAAGGGATGGAAGGCCTCCTTGGGACTCTAGTGCAGCTTCTGGGTTCCGATGATATAAATGTGGTCACCTGTGCAGCTGGAATTCTCTCTAACCTCACTT
GCAATAATTACAAAAACAAGATGATGGTGTGCCAAGTGGGTGGCATAGAGGCTCTTGTACGCACCGTCCTTCGTGCTGGTGACAGGGAAGACATCACTGAGCCTGCCATCTGTG
CTCTTCGTCATCTGACCAGCCGGCATCAGGAAGCCGAGATGGCCCAGAATGCCGTTCGCCTTCATTATGGACTGCCTGTTGTGGTTAAACTCCTGCACCCACCATCCCACTGGC
CTCTGATAAAGGCAACTGTTGGATTGATTCGAAACCTTGCCCTTTGCCCAGCAAATCATGCGCCTTTGCGGGAACAGGGTGCTATTCCACGACTAGTTCAGCTGCTTGTACGAG
CACATCAGGACACCCAACGGCGCACCTCCATGGGTGGAACGCAGCAGCAGTTTGTGGAGGGCGTGCGCATGGAGGAGATAGTAGAAGGGTGTACTGGAGCTCTCCACATCCTTG
CTCGGGACGTTCACAACCGGATTGTAATCCGAGGACTCAATACCATTCCATTGTTTGTGCAGTTGCTTTATTCTCCCATTGAAAATATCCAAAGAGTAGCTGCAGGGGTCCTCT
GTGAACTTGCTCAGGACAAGGAGGCTGCAGAGGCCATTGAAGCTGAGGGAGCCACAGCTCCCCTGACAGAGTTACTCCACTCCAGGAATGAAGGCGTGGCAACATACGCAGCTG
CTGTCCTATTCCGAATGTCTGAGGACAAGCCACAGGATTACAAGAAGCGGCTTTCAGTCGAGCTGACCAGTTCCCTCTTCAGGACAGAGCCAATGGCTTGGAATGAGACTGCAG
ATTAACCCGGGGAATTCCTCGAGAAGCTGGGGCTCGAGATCTAGAGTCGAGAAGCTTGATGATCTGCGCAGCACCATGGCCTGAAATAACCTCTGAAAGAGGAACTTGGTTAGG
TACCTTCTGAGGCGGAAAGAACCAGCTGTGGAATGTGTGTCAGTTAGGGTGTGGAAAGTCCCCAGGCTCCCCAGCAGGCAGAAGTATGCAAAGCATGCATCTCAATTAGTCAGC
AACCAGGTGTGGAAAGTCCCCAGGCTCCCCAGCAGGCAGAAGTATGCAAAGCATGCATCTCAATTAGTCAGCAACCATAGTCCCGCCCCTAACTCCGCCCATCCCGCCCCTAAC
TCCGCCCAGTTCCGCCCATTCTCCGCCCCATGGCTGACTAATTTTTTTTATTTATGCAGAGGCCGAGGCCGCCTCGGCCTCTGAGCTATTCCAGAAGTAGTGAGGAGGCTTTTT
TGGAGGCCTAGGCTTTTGCAAAAAGCTTACCATGACCGAGTACAAGCCCACGGTGCGCCTCGCCACCCGCGACGACGTCCCCAGGGCCGTACGCACCCTCGCCGCCGCGTTCGC
CGACTACCCCGCCACGCGCCACACCGTCGATCCGGACCGCCACATCGAGCGGGTCACCGAGCTGCAAGAACTCTTCCTCACGCGCGTCGGGCTCGACATCGGCAAGGTGTGGGT
CGCGGACGACGGCGCCGCGGTGGCGGTCTGGACCACGCCGGAGAGCGTCGAAGCGGGGGCGGTGTTCGCCGAGATCGGCCCGCGCATGGCCGAGTTGAGCGGTTCCCGGCTGGC
CGCGCAGCAACAGATGGAAGGCCTCCTGGCGCCGCACCGGCCCAAGGAGCCCGCGTGGTTCCTGGCCACCGTCGGCGTCTCGCCCGACCACCAGGGCAAGGGTCTGGGCAGCGC
CGTCGTGCTCCCCGGAGTGGAGGCGGCCGAGCGCGCCGGGGTGCCCGCCTTCCTGGAGACCTCCGCGCCCCGCAACCTCCCCTTCTACGAGCGGCTCGGCTTCACCGTCACCGC
CGACGTCGAGGTGCCCGAAGGACCGCGCGACCTGGTGCATGACCCGCAAGCCCGGTGCCTGACGCCCGCCCCACGACCCGCAGCGCCCGACCGAAAGGAGCGCACGACCCCATGC
CAGTCGACAATCAACCTCTGGATTACAAAATTTGTGAAAGATTGACTGGTATTCTTAACTATGTTGCTCCTTTTACGCTATGTGGATACGCTGCTTTAATGCCTTTGTATCATG
CTATTGCTTCCCGTATGGCTTTCATTTTCTCCTCCTTGTATAAATCCTGGTTGCTGTCTCTTTATGAGGAGTTGTGGCCCGTTGTCAGGCAACGTGGCGTGGTGTGCACTGTGT
TTGCTGACGCAACCCCCACTGGTTGGGGCATTGCCACCACCTGTCAGCTCCTTTCCGGGACTTTCGCTTTCCCCCTCCCTATTGCCACGGCGGAACTCATCGCCGCCTGCCTTG
CCCGCTGCTGGACAGGGGCTCGGCTGTTGGGCACTGACAATTCCGTGGTGTTGTCGGGGAAGCTGACGTCCTTTCCATGGCTGCTCGCCTGTGTTGCCACCTGGATTCTGCGCG
GGACGTCCTTCTGCTACGTCCCTTCGGCCCTCAATCCAGCGGACCTTCCTTCCCGCGGCCTGCTGCCGGCTCTGCGGCCTCTTCCGCGTCTTCGCCTTCGCCCTCAGACGAGTC
GGATCTCCCTTTGGGCCGCCTCCCCGCCTGGAATTCGAGCTCGGTACCTTTAAGACCAATGACTTACAAGGCAGCTGTAGATCTTAGCCACTTTTTAAAAGAAAAGGGGGGACT
GGAAGGGCTAATTCACTCCCAACGAAGACAAGATCTGCTTTTTGCTTGTACTGGGTCTCTCTGGTTAGACCAGATCTGAGCCTGGGAGCTCTCTGGCTAACTAGGGAACCCACT
GCTTAAGCCTCAATAAAGCTTGCCTTGAGTGCTTCAAGTAGTGTGTGCCCGTCTGTTGTGTGACTCTGGTAACTAGAGATCCCTCAGACCCTTTTAGTCAGTGTGGAAAATCTC
TAGCAGTAGTAGTTCATGTCATCTTATTATTCAGTATTTATAACTTGCAAAGAAATGAATATCAGAGAGTGAGAGGAACTTGTTTATTGCAGCTTATAATGGTTACAAATAAAG
CAATAGCATCACAAATTTCACAAATAAAGCATTTTTTTCACTGCATTCTAGTTGTGGTTTGTCCAAACTCATCAATGTATCTTATCATGTCTGGCTCTAGCTATCCCGCCCCTA
ACTCCGCCCAGTTCCGCCCATTCTCCGCCCCATGGCTGACTAATTTTTTTTATTTATGCAGAGGCCGAGGCCGCCTCGGCCTCTGAGCTATTCCAGAAGTAGTGAGGAGGCTTT
TTTGGAGGCCTAGGCTTTTGCGTCGAGACGTACCCAATTCGCCCTATAGTGAGTCGTATTACGCGCGCTCACTGGCCGTCGTTTTACAACGTCGTGACTGGGAAAACCCTGGCG

TTACCCAACTTAATCGCCTTGCAGCACATCCCCCTTTCGCCAGCTGGCGTAATAGCGAAGAGGCCCGCACCGATCGCCCTTCCCAACAGTTGCGCAGCCTGAATGGCGAATGGC
GCGACGCGCCCTGTAGCGGCGCATTAAGCGCGGCGGGTGTGGTGGTTACGCGCAGCGTGACCGCTACACTTGCCAGCGCCCTAGCGCCCGCTCCTTTCGCTTTCTTCCCTTCCT
TTCTCGCCACGTTCGCCGGCTTTCCCCGTCAAGCTCTAAATCGGGGGCTCCCTTTAGGGTTCCGATTTAGTGCTTTACGGCACCTCGACCCCAAAAAACTTGATTAGGGTGATG
GTTCACGTAGTGGGCCATCGCCCTGATAGACGGTTTTTCGCCCTTTGACGTTGGAGTCCACGTTCTTTAATAGTGGACTCTTGTTCCAAACTGGAACAACACTCAACCCTATCT
CGGTCTATTCTTTTGATTTATAAGGGATTTTGCCGATTTCGGCCTATTGGTTAAAAAATGAGCTGATTTAACAAAAATTTAACGCGAATTTTAACAAAATATTAACGTTTACAA
TTTCCCAGGTGGCACTTTTCGGGGAAATGTGCGCGGAACCCCTATTTGTTTATTTTTCTAAATACATTCAAATATGTATCCGCTCATGAGACAATAACCCTGATAAATGCTTCA
ATAATATTGAAAAAGGAAGAGTATGAGTATTCAACATTTCCGTGTCGCCCTTATTCCCTTTTTTGCGGCATTTTGCCTTCCTGTTTTTGCTCACCCAGAAACGCTGGTGAAAGT
AAAAGATGCTGAAGATCAGTTGGGTGCACGAGTGGGTTACATCGAACTGGATCTCAACAGCGGTAAGATCCTTGAGAGTTTTCGCCCCGAAGAACGTTTTCCAATGATGAGCAC
TTTTAAAGTTCTGCTATGTGGCGCGGTATTATCCCGTATTGACGCCGGGCAAGAGCAACTCGGTCGCCGCATACACTATTCTCAGAATGACTTGGTTGAGTACTCACCAGTCAC
AGAAAAGCATCTTACGGATGGCATGACAGTAAGAGAATTATGCAGTGCTGCCATAACCATGAGTGATAACACTGCGGCCAACTTACTTCTGACAACGATCGGAGGACCGAAGGA
GCTAACCGCTTTTTTGCACAACATGGGGGATCATGTAACTCGCCTTGATCGTTGGGAACCGGAGCTGAATGAAGCCATACCAAACGACGAGCGTGACACCACGATGCCTGTAGC
AATGGCAACAACGTTGCGCAAACTATTAACTGGCGAACTACTTACTCTAGCTTCCCGGCAACAATTAATAGACTGGATGGAGGCGGATAAAGTTGCAGGACCACTTCTGCGCTC
GGCCCTTCCGGCTGGCTGGTTTATTGCTGATAAATCTGGAGCCGGTGAGCGTGGGTCTCGCGGTATCATTGCAGCACTGGGGCCAGATGGTAAGCCCTCCCGTATCGTAGTTAT
CTACACGACGGGGAGTCAGGCAACTATGGATGAACGAAATAGACAGATCGCTGAGATAGGTGCCTCACTGATTAAGCATTGGTAACTGTCAGACCAAGTTTACTCATATATACT
TTAGATTGATTTAAAACTTCATTTTTAATTTAAAAGGATCTAGGTGAAGATCCTTTTTGATAATCTCATGACCAAAATCCCTTAACGTGAGTTTTCGTTCCACTGAGCGTCAGA
CCCCGTAGAAAAGATCAAAGGATCTTCTTGAGATCCTTTTTTTCTGCGCGTAATCTGCTGCTTGCAAACAAAAAAACCACCGCTACCAGCGGTGGTTTGTTTGCCGGATCAAGA
GCTACCAACTCTTTTTCCGAAGGTAACTGGCTTCAGCAGAGCGCAGATACCAAATACTGTCCTTCTAGTGTAGCCGTAGTTAGGCCACCACTTCAAGAACTCTGTAGCACCGCC
TACATACCTCGCTCTGCTAATCCTGTTACCAGTGGCTGCTGCCAGTGGCGATAAGTCGTGTCTTACCGGGTTGGACTCAAGACGATAGTTACCGGATAAGGCGCAGCGGTCGGG
CTGAACGGGGGGTTCGTGCACACAGCCCAGCTTGGAGCGAACGACCTACACCGAACTGAGATACCTACAGCGTGAGCTATGAGAAAGCGCCACGCTTCCCGAAGGGAGAAAGGC
GGACAGGTATCCGGTAAGCGGCAGGGTCGGAACAGGAGAGCGCACGAGGGAGCTTCCAGGGGGAAACGCCTGGTATCTTTATAGTCCTGTCGGGTTTCGCCACCTCTGACTTGA
GCGTCGATTTTTGTGATGCTCGTCAGGGGGGCGGAGCCTATGGAAAAACGCCAGCAACGCGGCCTTTTTACGGTTCCTGGCCTTTTGCTGGCCTTTTGCTCACATGTTCTTTCC
TGCGTTATCCCCTGATTCTGTGGATAACCGTATTACCGCCTTTGAGTGAGCTGATACCGCTCGCCGCAGCCGAACGACCGAGCGCAGCGAGTCAGTGAGCGAGGAAGCGGAAGA
GCGCCCAATACGCAAACCGCCTCTCCCCGCGCGTTGGCCGATTCATTAATGCAGCTGGCACGACAGGTTTCCCGACTGGAAAGCGGGCAGTGAGCGCAACGCAATTAATGTGAG
TTAGCTCACTCATTAGGCACCCCAGGCTTTACACTTTATGCTTCCGGCTCGTATGTTGTGTGGAATTGTGAGCGGATAACAATTTCACACAGGAAACAGCTATGACCATGATTA
CGCCAAGCGCGCAATTAACCCTCACTAAAGGGAACAAAAGCTGGAGCTGCAAGCTTAATGTAGTCTTATGCAATACTCTTGTAGTCTTGCAACATGGTAACGATGAGTTAGCAA
CATGCCTTACAAGGAGAGAAAAAGCACCGTGCATGCCGATTGGTGGAAGTAAGGTGGTACGATCGTGCCTTATTAGGAAGGCAACAGACGGGTCTGACATGGATTGGACGAACC
ACTGAATTGCCGCATTGCAGAGATATTGTATTTAAGTGCCTAGCTCGATACAATAAACGGGTCTCTCTGGTTAGACCAGATCTGAGCCTGGGAGCTCTCTGGCTAACTAGGGAA
CCCACTGCTTAAGCCTCAATAAAGCTTGCCTTGAGTGCTTCAAGTAGTGTGTGCCCGTCTGTTGTGTGACTCTGGTAACTAGAGATCCCTCAGACCCTTTTAGTCAGTGTGGAA
AATCTCTAGCAGTGGCGCCCGAACAGGGACCTGAAAGCGAAAGGGAAACCAGAGCTCTCTCGACGCAGGACTCGGCTTGCTGAAGCGCGCACGGCAAGAGGCGAGGGGCGGCGA
CTGGTGAGTACGCCAAAAATTTTGACTAGCGGAGGCTAGAAGGAGAGAGATGGGTGCGAGAGCGTCAGTATTAAGCGGGGGAGAATTAGATCGCGATGGGAAAAAATTCGGTTA
AGGCCAGGGGGAAAGAAAAAATATAAATTAAAACATATAGTATGGGCAAGCAGGGAGCTAGAACGATTCGCAGTTAATCCTGGCCTGTTAGAAACATCAGAAGGCTGTAGACAA
ATACTGGGACAGCTACAACCATCCCTTCAGACAGGATCAGAAGAACTTAGATCATTATATAATACAGTAGCAACCCTCTATTGTGTGCATCAAAGGATAGAGATAAAAGACACC
AAGGAAGCTTTAGACAAGATAGAGGAAGAGCAAAACAAAAGTAAGACCACCGCACAGCAAGCGGCCGCTGATCTTCAGACCTGGAGGAGGAGATATGAGGGACAATTGGAGAAG
TGAATTATATAAATATAAAGTAGTAAAAATTGAACCATTAGGAGTAGCACCCACCAAGGCAAAGAGAAGAGTGGTGCAGAGAGAAAAAAGAGCAGTGGGAATAGGAGCTTTGTT
CCTTGGGTTCTTGGGAGCAGCAGGAAGCACTATGGGCGCAGCCTCAATGACGCTGACGGTACAGGCCAGACAATTATTGTCTGGTATAGTGCAGCAGCAGAACAATTTGCTGAG
GGCTATTGAGGCGCAACAGCATCTGTTGCAACTCACAGTCTGGGGCATCAAGCAGCTCCAGGCAAGAATCCTGGCTGTGGAAAGATACCTAAAGGATCAACAGCTCCTGGGGAT
TTGGGGTTGCTCTGGAAAACTCATTTGCACCACTGCTGTGCCTTGGAATGCTAGTTGGAGTAATAAATCTCTGGAACAGATTTGGAATCACACGACCTGGATGGAGTGGGACAG
AGAAATTAACAATTACACAAGCTTAATACACTCCTTAATTGAAGAATCGCAAAACCAGCAAGAAAAGAATGAACAAGAATTATTGGAATTAGATAAATGGGCAAGTTTGTGGAA
TTGGTTTAACATAACAAATTGGCTGTGGTATATAAAATTATTCATAATGATAGTAGGAGGCTTGGTAGGTTTAAGAATAGTTTTTGCTGTACTTTCTATAGTGAATAGAGTTAG
GCAGGGATATTCACCATTATCGTTTCAGACCCACCTCCCAACCCCGAGGGGACCCGACAGGCCCGAAGGAATAGAAGAAGAAGGTGGAGAGAGAGACAGAGACAGATCCATTCG
ATTAGTGAACGGATCTCGACGGTATCGGTTAACTTTTAAAAGAAAAGGGGGGATTGGGGGGTACAGTGCAGGGGAAAGAATAGTAGACATAATAGCAACAGACATACAAACTAA
AGAATTACAAAAACAAATTACAAAAATTCAAAATTTTATCGATCACGAGACTAGCCTC

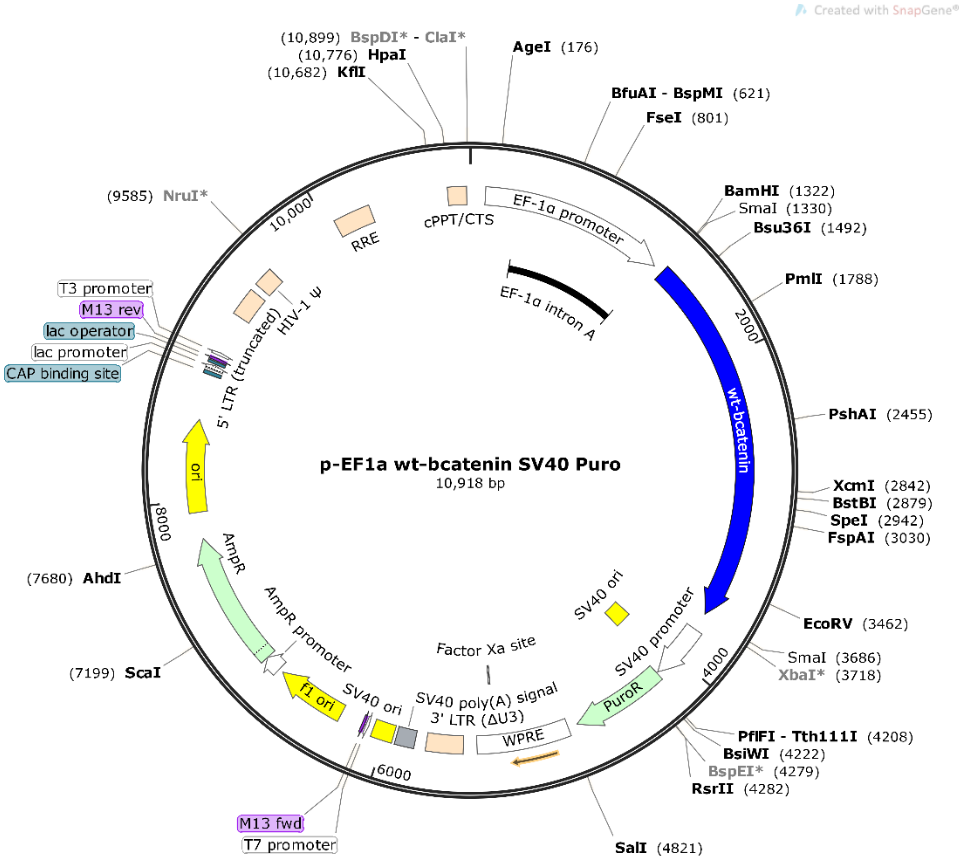

>pL-EF1a wt-βcatenin SV40 Puro

CTAGCCCCGATAAGCTTTGCAAAGATGGATAAAGTTTTAAACAGAGAGGAATCTTTGCAGCTAATGGACCTTCTAGGTCTTGAAAGGAGTGGGAATTGGCTCCGGTGCCCGTCA
GTGGGCAGAGCGCACATCGCCCACAGTCCCCGAGAAGTTGGGGGGAGGGGTCGGCAATTGAACCGGTGCCTAGAGAAGGTGGCGCGGGGTAAACTGGGAAAGTGATGTCGTGTA
CTGGCTCCGCCTTTTTCCCGAGGGTGGGGGAGAACCGTATATAAGTGCAGTAGTCGCCGTGAACGTTCTTTTTCGCAACGGGTTTGCCGCCAGAACACAGGTAAGTGCCGTGTG
TGGTTCCCGCGGGCCTGGCCTCTTTACGGGTTATGGCCCTTGCGTGCCTTGAATTACTTCCACTGGCTGCAGTACGTGATTCTTGATCCCGAGCTTCGGGTTGGAAGTGGGTGG
GAGAGTTCGAGGCCTTGCGCTTAAGGAGCCCCTTCGCCTCGTGCTTGAGTTGAGGCCTGGCCTGGGCGCTGGGGCCGCCGCGTGCGAATCTGGTGGCACCTTCGCGCCTGTCTC
GCTGCTTTCGATAAGTCTCTAGCCATTTAAAATTTTTGATGACCTGCTGCGACGCTTTTTTTCTGGCAAGATAGTCTTGTAAATGCGGGCCAAGATCTGCACACTGGTATTTCG
GTTTTTGGGGCCGCGGGCGGCGACGGGGCCCGTGCGTCCCAGCGCACATGTTCGGCGAGGCGGGGCCTGCGAGCGCGGCCACCGAGAATCGGACGGGGGTAGTCTCAAGCTGGC
CGGCCTGCTCTGGTGCCTGGCCTCGCGCCGCCGTGTATCGCCCCGCCCTGGGCGGCAAGGCTGGCCCGGTCGGCACCAGTTGCGTGAGCGGAAAGATGGCCGCTTCCCGGCCCT
GCTGCAGGGAGCTCAAAATGGAGGACGCGGCGCTCGGGAGAGCGGGCGGGTGAGTCACCCACACAAAGGAAAAGGGCCTTTCCGTCCTCAGCCGTCGCTTCATGTGACTCCACG
GAGTACCGGGCGCCGTCCAGGCACCTCGATTAGTTCTCGAGCTTTTGGAGTACGTCGTCTTTAGGTTGGGGGGAGGGGTTTTATGCGATGGAGTTTCCCCACACTGAGTGGGTG

GAGACTGAAGTTAGGCCAGCTTGGCACTTGATGTAATTCTCCTTGGAATTTGCCCTTTTTGAGTTTGGATCTTGGTTCATTCTCAAGCCTCAGACAGTGGTTCAAAGTTTTTTT
CTTCCATTTCAGGTGTCGTGAGGAATTCTGCAGTCGATCGACGGTACCGCGGGCCCTCCGCGGCCGCGGATCCCCCGGGCCACATGGCTACTCAAGCTGACCTGATGGAGTTGG
ACATGGCCATGGAGCCGGACAGAAAAGCTGCTGTCAGCCACTGGCAGCAGCAGTCTTACTTGGATTCTGGAATCCATTCTGGTGCCACCACCACAGCTCCTTCCCTGAGTGGCA
AGGGCAACCCTGAGGAAGAAGATGTTGACACCTCCCAAGTCCTTTATGAATGGGAGCAAGGCTTTTCCCAGTCCTTCACGCAAGAGCAAGTAGCTGATATTGACGGGCAGTATG
CAATGACTAGGGCTCAGAGGGTCCGAGCTGCCATGTTCCCTGAGACGCTAGATGAGGGCATGCAGATCCCATCCACGCAGTTTGACGCTGCTCATCCCACTAATGTCCAGCGCT
TGGCTGAACCATCACAGATGTTGAAACATGCAGTTGTCAATTTGATTAACTATCAGGATGACGCGGAACTTGCCACACGTGCAATTCCTGAGCTGACAAAACTGCTAAACGATG
AGGACCAGGTGGTAGTTAATAAAGCTGCTGTTATGGTCCATCAGCTTTCCAAAAAGGAAGCTTCCAGACATGCCATCATGCGCTCCCCTCAGATGGTGTCTGCCATTGTACGCA
CCATGCAGAATACAAATGATGTAGAGACAGCTCGTTGTACTGCTGGGACTCTGCACAACCTTTCTCACCACCGCGAGGGCTTGCTGGCCATCTTTAAGTCTGGTGGCATCCCAG
CGCTGGTGAAAATGCTTGGGTCACCAGTGGATTCTGTACTGTTCTACGCCATCACGACACTGCATAATCTCCTGCTCCATCAGGAAGGAGCTAAAATGGCAGTGCGCCTAGCTG
GTGGACTGCAGAAAATGGTTGCTTTGCTCAACAAAACAAACGTGAAATTCTTGGCTATTACAACAGACTGCCTTCAGATCTTAGCTTATGGCAATCAAGAGAGCAAGCTCATCA
TTCTGGCCAGTGGTGGACCCCAAGCCTTAGTAAACATAATGAGGACCTACACTTATGAGAAGCTTCTGTGGACCACAAGCAGAGTGCTGAAGGTGCTGTCTGTCTGCTCTAGCA
ACAAGCCGGCCATTGTAGAAGCTGGTGGGATGCAGGCACTGGGGCTTCATCTGACAGACCCAAGTCAGCGACTTGTTCAAAACTGTCTTTGGACTCTCAGAAACCTTTCAGATG
CAGCGACTAAGCAGGAAGGGATGGAAGGCCTCCTTGGGACTCTAGTGCAGCTTCTGGGTTCCGATGATATAAATGTGGTCACCTGTGCAGCTGGAATTCTCTCTAACCTCACTT
GCAATAATTACAAAAACAAGATGATGGTGTGCCAAGTGGGTGGCATAGAGGCTCTTGTACGCACCGTCCTTCGTGCTGGTGACAGGGAAGACATCACTGAGCCTGCCATCTGTG
CTCTTCGTCATCTGACCAGCCGGCATCAGGAAGCCGAGATGGCCCAGAATGCCGTTCGCCTTCATTATGGACTGCCTGTTGTGGTTAAACTCCTGCACCCACCATCCCACTGGC
CTCTGATAAAGGCAACTGTTGGATTGATTCGAAACCTTGCCCTTTGCCCAGCAAATCATGCGCCTTTGCGGGAACAGGGTGCTATTCCACGACTAGTTCAGCTGCTTGTACGAG
CACATCAGGACACCCAACGGCGCACCTCCATGGGTGGAACGCAGCAGCAGTTTGTGGAGGGCGTGCGCATGGAGGAGATAGTAGAAGGGTGTACTGGAGCTCTCCACATCCTTG
CTCGGGACGTTCACAACCGGATTGTAATCCGAGGACTCAATACCATTCCATTGTTTGTGCAGTTGCTTTATTCTCCCATTGAAAATATCCAAAGAGTAGCTGCAGGGGTCCTCT
GTGAACTTGCTCAGGACAAGGAGGCTGCAGAGGCCATTGAAGCTGAGGGAGCCACAGCTCCCCTGACAGAGTTACTCCACTCCAGGAATGAAGGCGTGGCAACATACGCAGCTG
CTGTCCTATTCCGAATGTCTGAGGACAAGCCACAGGATTACAAGAAGCGGCTTTCAGTCGAGCTGACCAGTTCCCTCTTCAGGACAGAGCCAATGGCTTGGAATGAGACTGCAG
ATCTTGGACTGGACATTGGTGCCCAGGGAGAAGCCCTTGGATATCGCCAGGATGATCCCAGCTACCGTTCTTTTCACTCTGGTGGATACGGCCAGGATGCCTTGGGGATGGACC
CTATGATGGAGCATGAGATGGGTGGCCACCACCCTGGTGCTGACTATCCAGTTGATGGGCTGCCTGATCTGGGACACGCCCAGGACCTCATGGATGGGCTGCCCCCAGGTGATA
GCAATCAGCTGGCCTGGTTTGATACTGACCTGTAACCCGGGGAATTCCTCGAGAAGCTGGGGCTCGAGATCTAGAGTCGAGAAGCTTGATGATCTGCGCAGCACCATGGCCTGA
AATAACCTCTGAAAGAGGAACTTGGTTAGGTACCTTCTGAGGCGGAAAGAACCAGCTGTGGAATGTGTGTCAGTTAGGGTGTGGAAAGTCCCCAGGCTCCCCAGCAGGCAGAAG
TATGCAAAGCATGCATCTCAATTAGTCAGCAACCAGGTGTGGAAAGTCCCCAGGCTCCCCAGCAGGCAGAAGTATGCAAAGCATGCATCTCAATTAGTCAGCAACCATAGTCCC
GCCCCTAACTCCGCCCATCCCGCCCCTAACTCCGCCCAGTTCCGCCCATTCTCCGCCCCATGGCTGACTAATTTTTTTTATTTATGCAGAGGCCGAGGCCGCCTCGGCCTCTGA
GCTATTCCAGAAGTAGTGAGGAGGCTTTTTTGGAGGCCTAGGCTTTTGCAAAAAGCTTACCATGACCGAGTACAAGCCCACGGTGCGCCTCGCCACCCGCGACGACGTCCCCAG
GGCCGTACGCACCCTCGCCGCCGCGTTCGCCGACTACCCCGCCACGCGCCACACCGTCGATCCGGACCGCCACATCGAGCGGGTCACCGAGCTGCAAGAACTCTTCCTCACGCG
CGTCGGGCTCGACATCGGCAAGGTGTGGGTCGCGGACGACGGCGCCGCGGTGGCGGTCTGGACCACGCCGGAGAGCGTCGAAGCGGGGGCGGTGTTCGCCGAGATCGGCCCGCG
CATGGCCGAGTTGAGCGGTTCCCGGCTGGCCGCGCAGCAACAGATGGAAGGCCTCCTGGCGCCGCACCGGCCCAAGGAGCCCGCGTGGTTCCTGGCCACCGTCGGCGTCTCGCC
CGACCACCAGGGCAAGGGTCTGGGCAGCGCCGTCGTGCTCCCCGGAGTGGAGGCGGCCGAGCGCGCCGGGGTGCCCGCCTTCCTGGAGACCTCCGCGCCCCGCAACCTCCCCTT
CTACGAGCGGCTCGGCTTCACCGTCACCGCCGACGTCGAGTGCCCGAAGGACCGCGCGACCTGGTGCATGACCCGCAAGCCCGGTGCCTGACGCCCGCCCCACGACCCGCAGCG
CCCGACCGAAAGGAGCGCACGACCCCATGCCAGTCGACAATCAACCTCTGGATTACAAAATTTGTGAAAGATTGACTGGTATTCTTAACTATGTTGCTCCTTTTACGCTATGTG
GATACGCTGCTTTAATGCCTTTGTATCATGCTATTGCTTCCCGTATGGCTTTCATTTTCTCCTCCTTGTATAAATCCTGGTTGCTGTCTCTTTATGAGGAGTTGTGGCCCGTTG
TCAGGCAACGTGGCGTGGTGTGCACTGTGTTTGCTGACGCAACCCCCACTGGTTGGGGCATTGCCACCACCTGTCAGCTCCTTTCCGGGACTTTCGCTTTCCCCCTCCCTATTG
CCACGGCGGAACTCATCGCCGCCTGCCTTGCCCGCTGCTGGACAGGGGCTCGGCTGTTGGGCACTGACAATTCCGTGGTGTTGTCGGGGAAGCTGACGTCCTTTCCATGGCTGC
TCGCCTGTGTTGCCACCTGGATTCTGCGCGGGACGTCCTTCTGCTACGTCCCTTCGGCCCTCAATCCAGCGGACCTTCCTTCCCGCGGCCTGCTGCCGGCTCTGCGGCCTCTTC
CGCGTCTTCGCCTTCGCCCTCAGACGAGTCGGATCTCCCTTTGGGCCGCCTCCCCGCCTGGAATTCGAGCTCGGTACCTTTAAGACCAATGACTTACAAGGCAGCTGTAGATCT
TAGCCACTTTTTAAAAGAAAAGGGGGGACTGGAAGGGCTAATTCACTCCCAACGAAGACAAGATCTGCTTTTTGCTTGTACTGGGTCTCTCTGGTTAGACCAGATCTGAGCCTG
GGAGCTCTCTGGCTAACTAGGGAACCCACTGCTTAAGCCTCAATAAAGCTTGCCTTGAGTGCTTCAAGTAGTGTGTGCCCGTCTGTTGTGTGACTCTGGTAACTAGAGATCCCT
CAGACCCTTTTAGTCAGTGTGGAAAATCTCTAGCAGTAGTAGTTCATGTCATCTTATTATTCAGTATTTATAACTTGCAAAGAAATGAATATCAGAGAGTGAGAGGAACTTGTT
TATTGCAGCTTATAATGGTTACAAATAAAGCAATAGCATCACAAATTTCACAAATAAAGCATTTTTTTCACTGCATTCTAGTTGTGGTTTGTCCAAACTCATCAATGTATCTTA
TCATGTCTGGCTCTAGCTATCCCGCCCCTAACTCCGCCCAGTTCCGCCCATTCTCCGCCCCATGGCTGACTAATTTTTTTTATTTATGCAGAGGCCGAGGCCGCCTCGGCCTCT
GAGCTATTCCAGAAGTAGTGAGGAGGCTTTTTTGGAGGCCTAGGCTTTTGCGTCGAGACGTACCCAATTCGCCCTATAGTGAGTCGTATTACGCGCGCTCACTGGCCGTCGTTT
TACAACGTCGTGACTGGGAAAACCCTGGCGTTACCCAACTTAATCGCCTTGCAGCACATCCCCCTTTCGCCAGCTGGCGTAATAGCGAAGAGGCCCGCACCGATCGCCCTTCCC
AACAGTTGCGCAGCCTGAATGGCGAATGGCGCGACGCGCCCTGTAGCGGCGCATTAAGCGCGGCGGGTGTGGTGGTTACGCGCAGCGTGACCGCTACACTTGCCAGCGCCCTAG
CGCCCGCTCCTTTCGCTTTCTTCCCTTCCTTTCTCGCCACGTTCGCCGGCTTTCCCCGTCAAGCTCTAAATCGGGGGCTCCCTTTAGGGTTCCGATTTAGTGCTTTACGGCACC
TCGACCCCAAAAAACTTGATTAGGGTGATGGTTCACGTAGTGGGCCATCGCCCTGATAGACGGTTTTTCGCCCTTTGACGTTGGAGTCCACGTTCTTTAATAGTGGACTCTTGT
TCCAAACTGGAACAACACTCAACCCTATCTCGGTCTATTCTTTTGATTTATAAGGGATTTTGCCGATTTCGGCCTATTGGTTAAAAAATGAGCTGATTTAACAAAAATTTAACG
CGAATTTTAACAAAATATTAACGTTTACAATTTCCCAGGTGGCACTTTTCGGGGAAATGTGCGCGGAACCCCTATTTGTTTATTTTTCTAAATACATTCAAATATGTATCCGCT
CATGAGACAATAACCCTGATAAATGCTTCAATAATATTGAAAAAGGAAGAGTATGAGTATTCAACATTTCCGTGTCGCCCTTATTCCCTTTTTTGCGGCATTTTGCCTTCCTGT
TTTTGCTCACCCAGAAACGCTGGTGAAAGTAAAAGATGCTGAAGATCAGTTGGGTGCACGAGTGGGTTACATCGAACTGGATCTCAACAGCGGTAAGATCCTTGAGAGTTTTCG
CCCCGAAGAACGTTTTCCAATGATGAGCACTTTTAAAGTTCTGCTATGTGGCGCGGTATTATCCCGTATTGACGCCGGGCAAGAGCAACTCGGTCGCCGCATACACTATTCTCA
GAATGACTTGGTTGAGTACTCACCAGTCACAGAAAAGCATCTTACGGATGGCATGACAGTAAGAGAATTATGCAGTGCTGCCATAACCATGAGTGATAACACTGCGGCCAACTT
ACTTCTGACAACGATCGGAGGACCGAAGGAGCTAACCGCTTTTTTGCACAACATGGGGGATCATGTAACTCGCCTTGATCGTTGGGAACCGGAGCTGAATGAAGCCATACCAAA
CGACGAGCGTGACACCACGATGCCTGTAGCAATGGCAACAACGTTGCGCAAACTATTAACTGGCGAACTACTTACTCTAGCTTCCCGGCAACAATTAATAGACTGGATGGAGGC
GGATAAAGTTGCAGGACCACTTCTGCGCTCGGCCCTTCCGGCTGGCTGGTTTATTGCTGATAAATCTGGAGCCGGTGAGCGTGGGTCTCGCGGTATCATTGCAGCACTGGGGCC
AGATGGTAAGCCCTCCCGTATCGTAGTTATCTACACGACGGGGAGTCAGGCAACTATGGATGAACGAAATAGACAGATCGCTGAGATAGGTGCCTCACTGATTAAGCATTGGTA
ACTGTCAGACCAAGTTTACTCATATATACTTTAGATTGATTTAAAACTTCATTTTTAATTTAAAAGGATCTAGGTGAAGATCCTTTTTGATAATCTCATGACCAAAATCCCTTA
ACGTGAGTTTTCGTTCCACTGAGCGTCAGACCCCGTAGAAAAGATCAAAGGATCTTCTTGAGATCCTTTTTTTCTGCGCGTAATCTGCTGCTTGCAAACAAAAAAACCACCGCT
ACCAGCGGTGGTTTGTTTGCCGGATCAAGAGCTACCAACTCTTTTTCCGAAGGTAACTGGCTTCAGCAGAGCGCAGATACCAAATACTGTCCTTCTAGTGTAGCCGTAGTTAGG
CCACCACTTCAAGAACTCTGTAGCACCGCCTACATACCTCGCTCTGCTAATCCTGTTACCAGTGGCTGCTGCCAGTGGCGATAAGTCGTGTCTTACCGGGTTGGACTCAAGACG
ATAGTTACCGGATAAGGCGCAGCGGTCGGGCTGAACGGGGGGTTCGTGCACACAGCCCAGCTTGGAGCGAACGACCTACACCGAACTGAGATACCTACAGCGTGAGCTATGAGA
AAGCGCCACGCTTCCCGAAGGGAGAAAGGCGGACAGGTATCCGGTAAGCGGCAGGGTCGGAACAGGAGAGCGCACGAGGGAGCTTCCAGGGGGAAACGCCTGGTATCTTTATAG
TCCTGTCGGGTTTCGCCACCTCTGACTTGAGCGTCGATTTTTGTGATGCTCGTCAGGGGGGCGGAGCCTATGGAAAAACGCCAGCAACGCGGCCTTTTTACGGTTCCTGGCCTT
TTGCTGGCCTTTTGCTCACATGTTCTTTCCTGCGTTATCCCCTGATTCTGTGGATAACCGTATTACCGCCTTTGAGTGAGCTGATACCGCTCGCCGCAGCCGAACGACCGAGCG
CAGCGAGTCAGTGAGCGAGGAAGCGGAAGAGCGCCCAATACGCAAACCGCCTCTCCCCGCGCGTTGGCCGATTCATTAATGCAGCTGGCACGACAGGTTTCCCGACTGGAAAGC
GGGCAGTGAGCGCAACGCAATTAATGTGAGTTAGCTCACTCATTAGGCACCCCAGGCTTTACACTTTATGCTTCCGGCTCGTATGTTGTGTGGAATTGTGAGCGGATAACAATT
TCACACAGGAAACAGCTATGACCATGATTACGCCAAGCGCGCAATTAACCCTCACTAAAGGGAACAAAAGCTGGAGCTGCAAGCTTAATGTAGTCTTATGCAATACTCTTGTAG
TCTTGCAACATGGTAACGATGAGTTAGCAACATGCCTTACAAGGAGAGAAAAAGCACCGTGCATGCCGATTGGTGGAAGTAAGGTGGTACGATCGTGCCTTATTAGGAAGGCAA
CAGACGGGTCTGACATGGATTGGACGAACCACTGAATTGCCGCATTGCAGAGATATTGTATTTAAGTGCCTAGCTCGATACAATAAACGGGTCTCTCTGGTTAGACCAGATCTG
AGCCTGGGAGCTCTCTGGCTAACTAGGGAACCCACTGCTTAAGCCTCAATAAAGCTTGCCTTGAGTGCTTCAAGTAGTGTGTGCCCGTCTGTTGTGTGACTCTGGTAACTAGAG
ATCCCTCAGACCCTTTTAGTCAGTGTGGAAAATCTCTAGCAGTGGCGCCCGAACAGGGACCTGAAAGCGAAAGGGAAACCAGAGCTCTCTCGACGCAGGACTCGGCTTGCTGAA
GCGCGCACGGCAAGAGGCGAGGGGCGGCGACTGGTGAGTACGCCAAAAATTTTGACTAGCGGAGGCTAGAAGGAGAGAGATGGGTGCGAGAGCGTCAGTATTAAGCGGGGGAGA
ATTAGATCGCGATGGGAAAAAATTCGGTTAAGGCCAGGGGGAAAGAAAAAATATAAATTAAAACATATAGTATGGGCAAGCAGGGAGCTAGAACGATTCGCAGTTAATCCTGGC
CTGTTAGAAACATCAGAAGGCTGTAGACAAATACTGGGACAGCTACAACCATCCCTTCAGACAGGATCAGAAGAACTTAGATCATTATATAATACAGTAGCAACCCTCTATTGT
GTGCATCAAAGGATAGAGATAAAAGACACCAAGGAAGCTTTAGACAAGATAGAGGAAGAGCAAAACAAAAGTAAGACCACCGCACAGCAAGCGGCCGCTGATCTTCAGACCTGG
AGGAGGAGATATGAGGGACAATTGGAGAAGTGAATTATATAAATATAAAGTAGTAAAAATTGAACCATTAGGAGTAGCACCCACCAAGGCAAAGAGAAGAGTGGTGCAGAGAGA
AAAAAGAGCAGTGGGAATAGGAGCTTTGTTCCTTGGGTTCTTGGGAGCAGCAGGAAGCACTATGGGCGCAGCCTCAATGACGCTGACGGTACAGGCCAGACAATTATTGTCTGG
TATAGTGCAGCAGCAGAACAATTTGCTGAGGGCTATTGAGGCGCAACAGCATCTGTTGCAACTCACAGTCTGGGGCATCAAGCAGCTCCAGGCAAGAATCCTGGCTGTGGAAAG
ATACCTAAAGGATCAACAGCTCCTGGGGATTTGGGGTTGCTCTGGAAAACTCATTTGCACCACTGCTGTGCCTTGGAATGCTAGTTGGAGTAATAAATCTCTGGAACAGATTTG
GAATCACACGACCTGGATGGAGTGGGACAGAGAAATTAACAATTACACAAGCTTAATACACTCCTTAATTGAAGAATCGCAAAACCAGCAAGAAAAGAATGAACAAGAATTATT
GGAATTAGATAAATGGGCAAGTTTGTGGAATTGGTTTAACATAACAAATTGGCTGTGGTATATAAAATTATTCATAATGATAGTAGGAGGCTTGGTAGGTTTAAGAATAGTTTT
TGCTGTACTTTCTATAGTGAATAGAGTTAGGCAGGGATATTCACCATTATCGTTTCAGACCCACCTCCCAACCCCGAGGGGACCCGACAGGCCCGAAGGAATAGAAGAAGAAGG
TGGAGAGAGAGACAGAGACAGATCCATTCGATTAGTGAACGGATCTCGACGGTATCGGTTAACTTTTAAAAGAAAAGGGGGGATTGGGGGGTACAGTGCAGGGGAAAGAATAGT
AGACATAATAGCAACAGACATACAAACTAAAGAATTACAAAAACAAATTACAAAAATTCAAAATTTTATCGATCACGAGACTAGCCTC

#### SI3 - sgRNA cloning oligonucleotides

| sgRNA | Distance from ATG (base pairs) | Top oligo sequence (5'-3') Target sequence in bold |
|---|---|---|
| sgRNA-1 | -9 (Intron 1\|2 Exon2) | CACCGGCGTGGACAATGGCTACTCA |
| sgRNA-2 | +309 (Exon 3) | CACCGATGGAGTTGGACATGGCCA |
| sgRNA-3 | +5265 (Exon 9) | CACCGTACGCACCGTCCTTCGTGC |
| sgRNA-4 | - 116 (Intron 1\|2) | CACCGTAGCAGAATCACGGTGACC |
| sgRNA-5 | +9572 (Exon 15) | CACCGTCTGAACGTGCATTGTGAT |

List of top oligonucleotides used for cloning sgRNAs into px459-spCas9-Puro digested with BbsI.

#### SI4 - short hairpin cloning oligonucleotides

| **Short hairpin** | **ENSEMBL Gene_ID** | **Top oligo sequence (5'-3')** |
|---|---|---|
| **sh-ßcat1 (CDS)** | **ENSMUSG 00000006932** | CCGG**TCTAACCTCACTTGCAATAAT**CTCGAGATTATTGCAAGTGAGGTTAGATTTTT |
| **sh-ßcat2 (CDS)** | **ENSMUSG 00000006932** | CCGG**GCTGATATTGACGGGCAGTAT**CTCGAGATACTGCCCGTCAATA |
| **sh-ßcat3 (3'UTR)** | **ENSMUSG** | CCGG**GGCGTTATCAAACCCTAGCCTT**CTCGAGA |
| **sh-Tcf1** | **ENSMUSG 00000000782** | CCGG**GTTCACCCACCCATCCTTGAT**CTCGAGATCAAGGATGGGTGGGTGAACTTTTT |
| **sh-Lef1** | **ENSMUSG 00000027985** | CCGG**TGGTCAGCGCGAGACAATTAT**CTCGAGATAATTGTCTCGCGCTGACCATTTTTG |
| **sh-Control** | | CCGG**GTCACGATAAGACAATGAT**CTCGAGATCATTGTCTTATCGTGACTTTTT |

List of top oligonucleotides used for cloning short hairpins into pLKO Hygro digested with AgeI/EcoRI. Target sequences are in bold.

#### SI4 - Ctnnb1 deletions and genotyping assays

| **sgRNAs Combination** | Expected deletion | **PCR genotype assay:** |
|---|---|---|
| **sgRNA-1+ sgRNA-3** | **4970 bp** | **Forward 1:** GAATCACGGTGACCTGGGTT<br>**Reverse 1:** GACCCTCTGAGCCCTAGTCA<br>**Reverse 2:**CAGTTCACCTTTATCAGAGGCCAG<br><br>**Wild-type product: F1+R1 -> 824 bp**<br>**Deletion product: F1+R2 -> 595 bp***<br><br>***Wild-type product of F1+R2 is 5565 bp, not amplified thanks to reduced extension timing.** |
| **sgRNA-1+ sgRNA-3** | **5287 bp** | **Forward 1:** GAATCACGGTGACCTGGGTT<br>**Reverse 1:** GACCCTCTGAGCCCTAGTCA<br>**Reverse 2:** CAGTTCACCTTTATCAGAGGCCAG<br><br>**Wild-type product: F1+R1 -> 824 bp**<br>**Deletion product: F1+R2 -> 278 bp**<br><br>***Wild-type product of F1+R2 is 5565 bp, not amplified thanks to reduced extension timing.** |
| **sgRNA-4 + sgRNA-5** | **9701 bp** | **Forward 1:** CACCGTATGCCTACAATCTGTTTCTA<br>**Reverse 1:** GACCCTCTGAGCCCTAGTCA<br>**Reverse 2:** CTACACAATGTTACACGTCTCCAGAT<br><br>**Wild-type product: F1+R1-> 951 bp**<br>**Deletion product: F1+R2 -> 551**<br><br>***Wild-type product of F1+R2 is 10252 bp, not amplified thanks to reduced extension timing.** |

#### SI5 – Antibodies

| Antibody target | Raised in | Working dilution | Supplier and catalogue number |
|---|---|---|---|
| **β-catenin (anti C-terminal)** | **Mouse** | **1:500 /1:1000 (WB)<br>1:100/1:500 (IF)** | **BD (#MAB-318)** |
| **ß-catenin (anti-N-terminal)** | **Rabbit** | **1:500 (IF)/1:1000(WB)** | **Cell Signaling (#9581)** |
| **Oct4** | **Mouse** | **1:100/1:200 (IF)<br>1:500/1:1000 (WB)** | **Santa Cruz (sc-5279)** |
| **Tubulin** | **Mouse** | **1:1000/1:2000 (WB)** | **Sigma (T0198)** |
| **Nanog** | **Rabbit** | **1:300 (IF)/1:1000 (WB)** | **Calbiochem (#SC1000)** |
| **Plakoglobin** | **Mouse** | **1:100 (IF)/1:1000 (WB)** | **BD (610253)** |
| **E-cadherin** | **Mouse** | **1:100(IF)/1:1000(WB)** | **BD (#610182)** |
| **E-cadherin Alexa-647 conj.** | **Rat** | **0.5 µg x 10^6 cells(FACS)** | **Biolegend (#147308)** |
| **Sox2** | **Rabbit** | **1:1000 (WB)** | **Cell signaling (#3579)** |

List of primary antibodies used for western blot (WB), immunofluorescence (IF) or flow cytometry (FACS) experiments. Anti C-terminal **β**-catenin primary antibody was used in all the western blot and immunofluorescences of this study to probe for **β**-catenin expresión.

Anti N-terminal **β**-catenin was additonally used in Figure 5b and 5c.

#### SI6 – List of qRT-PCR oligonucleotides

| **Target** | Forward (5'-3') | Reverse (5'-3') |
|---|---|---|
| **Axin2** | GAGAGTGAGCGGCAGAGC | CGGCTGACTCGTTCTCCT |
| **Ctnnb1#1** | CGACACTGCATAATCTCCTGCTCC | GGTCCACCACTGGCCAGAATGAT |
| **Ctnnb1#2** | GTGGAAGTTTCTCACGTTGATGTT | ACAGCTGTATAGAGAGAAAGGCTG |
| **GAPDH** | GTATGACTCCACTCACGGCAAA | TTCCATTCTCGGCCTTG |
| **Tcf7 (Tcf1)** | GCTGCCTGAGGTCAGAGAAT | CCCCAGCTTTCTCCACTCTA |
| **Lef1** | CAGCCCGTGAGAAGGCTA | CTGGAGGATCGCACAGAGA |
| **Sp5** | CCCTCCAGACTTTTCCACCC | GGCTGCAGGTGTGTTTTCTG |
| **Cdx1** | TCTACACAGACCACCAACGC | TGCGCCGGATAGTGATGTAC |

List of oligonucleotides used for qRT-PCR assays.
